# MicroRNA miR-92b-3p regulation of a cardiovascular gene regulatory network in the developing branchial arches

**DOI:** 10.1101/2024.07.28.605500

**Authors:** Sian Goldsworthy, Marta Losa, Nicoletta Bobola, Sam Griffiths-Jones

**Affiliations:** Faculty of Biology, Medicine and Health, University of Manchester, UK

## Abstract

Vertebrate branchial arches (BAs) are a developmental paradigm, undergoing coordinated differentiation and morphogenesis to form various adult derivative tissues. MicroRNAs can strengthen gene regulatory networks (GRNs) to promote developmental stability. To interrogate the contribution of microRNAs to BA development, we generated a novel microRNA-sequencing dataset from mouse BAs. We identified 550 expressed microRNAs, of which approximately 20% demonstrate significant differential expression across BA domains. The three most posterior BAs and the connecting outflow tract (PBA/OFT) are enriched in biological processes linked to cardiovascular development. We identified enriched predicted microRNA-target interactions with PBA/OFT upregulated cardiovascular genes and validated transcripts encoding for two fundamental cardiac transcription factors (TFs), *Gata6* and *Tbx20,* as targets of miR-92b-3p. Furthermore, we demonstrated that miR-92b-3p can downregulate endogenous *GATA6* and *TBX20* in human embryonic stem cells (hESCs) undergoing cardiomyocyte differentiation, consistent with conservation of these microRNA-target interactions in a cardiogenic setting. miR-92b-3p has previously been shown to target two other cardiac TFs, *Hand2* and *Mef2D.* Therefore, we hypothesise that miR-92b-3p acts to stabilise cardiovascular GRNs during PBA/OFT development, through acting in multiple microRNA-mediated coherent feedforward loops.

## 1. INTRODUCTION

Vertebrate branchial arches (BAs) represent a principal developmental model that incorporates segmented design, cell migration and tissue specification. The BA transient domains arise during mid-embryonic development and comprise a series of outgrowths on each side of the embryonic head and pharynx, ultimately contributing to mature head, neck, and cardiovascular structures (1). Correct formation of these mature structures relies on the interplay between distinct embryonic populations found within the BAs: a mesenchymal core containing mesoderm and cranial neural crest (NC) surrounded by endoderm and ectoderm epithelia (1,2). Cranial NC cells that populate the BAs, originate from the hindbrain, and undergo epithelial to mesenchymal transition to migrate in discrete streams to the BAs (3). These cells express distinct members of *Hox* cluster genes, which is central to BA patterning (1). As cranial NC have the potential to give rise to a range of tissues including muscular, skeletal, vascular, and nervous, failure of migration and differentiation has been associated with multiple congenital defects (4).

In mammals there are five pairs of BAs: BA1, BA2, BA3, BA4, and BA6 (1). In this study we refer to the latter three, and the connected outflow tract (OFT), as the PBA/OFT. The PBA/OFT domain gives rise to multiple structures of the cardiovascular system, partly due to the migration of a subpopulation of cranial NC called cardiac NC. This subpopulation migrates from more caudal regions of the hindbrain, compared to other cranial NC, into the PBA/OFT (5–7). Cardiac NC initially form smooth muscle cells for the BA arterial system, connecting the embryonic heart to the dorsal aorta (8). BA arteries undergo subsequent remodelling into the great arteries, with cardiac NC contributing to the septation of the OFT into the aorta and pulmonary artery (9,10). Additional cardiovascular derivatives of cardiac NC cells include parasympathetic nerves and the surrounding cells of the His-Purkinje system (11), as well as a reported small proportion of cardiomyocytes (12). The PBA also contribute to the formation of carotid arteries supplying blood to the head and neck (1).

MicroRNAs are important factors in regulating embryonic development, with multiple studies validating their roles in cell fate decision and tissue patterning (13–16). Generally, microRNAs are understood to execute two different mechanisms of regulation. The first mechanism regards microRNAs as “on-off” switches, whereby they are anticorrelated in expression with their targets; this mechanism has been associated with earlier embryonic development (17). The second mode sees microRNAs as fine-tuners of their targets, stabilising their expression and attenuating noise brought about by stochasticity. These microRNAs are reported to be expressed later during development, and generally show overall weaker repressive abilities (17). However, weaker repression across many targets has been shown to stabilise GRNs through cumulative effects (18–20).

MicroRNA-mediated regulation extends to cardiovascular and craniofacial development. Disruption of the microRNA biogenesis pathway, through NC conditional *Dicer* loss of function, has been implicated in abnormal BA vessel remodelling (21) and OFT morphogenesis (22). Furthermore, *Dicer* loss of function in mouse models resembles several congenital phenotypes observed in DiGeorge Syndrome (DS) patients (21). DS individuals commonly have a genomic deletion encoding *DGCR8* (23), which is essential for microRNA processing (24). In addition to incorrect BA arterial remodelling, disrupting microRNA biogenesis also leads to craniofacial defects, due to aberrant skeletal formation and muscular maldevelopment (21). Taken together, microRNA biogenesis and expression are important for normal development of BA derivatives.

In this study we present novel microRNA-seq datasets, characterising global microRNA expression during mouse BA development. We find 550 microRNAs expressed across BA1, BA2, and the PBA/OFT, with the number of upregulated microRNAs progressively increasing across the anterior-posterior axis. Using corresponding BA RNA-seq datasets (25), we identified candidate microRNA target genes enriched for biological processes linked to cardiovascular development. Using *in silico* microRNA target prediction and *in vitro* microRNA target validation, we identify a role for miR-92b-3p as a key cardiovascular developmental regulator, adding to previous studies that have demonstrated miR-92b-3p regulates the cardiac TFs *Hand2* and *Mef2d* (26–28). We hypothesise that miR-92b-3p works in multiple microRNA-mediated coherent feedforward loops within cardiovascular GRNs, ultimately stabilising target gene expression during mammalian PBA/OFT development.

## 2. MATERIALS AND METHODS

### 2.1 BA dissection, RNA extraction and library preparation

Wild type (CD1) mice were time-mated to obtain tissue for microdissection. Animal experiments followed local legislations regarding housing, husbandry, and welfare (ASPA 1986, UK). Embryos were collected at E10.5 and E11.5 and accurately staged by counting somites. BA1, BA2 and PBA/OFT tissues were dissected and snap frozen on dry ice, and stored at -80°C until RNA extraction. The OFT was harvested with the PBA as it acts as a landmark feature during dissection and maintains physical integrity of the PBA. BAs from three closely staged embryos were pooled for each library, with BA1, BA2 and PBA/OFT from the same embryos being used for each timepoint replicate. Total RNA was extracted using the miRNeasy micro kit (Qiagen, #217084) following the manufacturer’s instructions, eluted into RNase-free water and stored at -20°C until library preparation. Small RNA libraries were generated using the NEBNext Small RNA Library Prep Set (New England BioLabs, #E7330S) following manufacturer’s instructions. For size selection we used gel separation, and extracted amplified microRNA cDNA bands corresponding to 140bp. NEBNext Index primer sequences and respective libraries are listed in Supplementary Table 1. Libraries were quality checked using the Agilent 2200 BioAnalyzer TapeStation and sequenced on the Illumina HiSeq 4000 at The University of Manchester Genomics Technologies Core Facility. Small RNA-seq libraries have been deposited under project accession PRJEB64007 available from the European Nucleotide Archive.

### 2.2 Small RNA-seq analysis

To remove NEBNext Small RNA adapter sequences from our libraries we used cutadapt v1.8 (29). Adapter-trimmed reads were filtered to keep those 18-25nt in length, and mapped against mouse tRNAs (mm10, GtRNAdb v18.1 (30)) and rRNAs (*Mus musculus*, Silva SSU/LSU r138.1 (31,32)), using Bowtie v1.1.0 (33). Reads that mapped to tRNAs/rRNAs were discarded from further analysis. Remaining reads were mapped to the mm10 primary fasta file (GRCm38, release M23) using bowtie v1.1.0 (33) with the following settings: bowtie -v1 -a -m5 –best –strata. To predict novel microRNAs we used miRDeep2 v0.1.3 (34) with combined filtered reads from all our small RNA-seq libraries. Reference microRNAs included stem-loop mouse, mature mouse and mature rat microRNAs, all downloaded from miRBase v22 (35). Novel pre-microRNAs were filtered using the following criteria: ≥30 0-mismatch reads for the mature arm and ≥10 0-mismatch reads for the star arm, no internal sub-hairpins, ≥50% 5’ arm homogeneity, 0-4nt overhang at the 3’ arm, hairpin free energy ≤ -0.2 kcal/mol/nt. Novel pre-microRNAs were filtered using custom Python scripts available at github.com/SianGol. Pre-microRNA sequences that met all of the above criteria were input into Rfam v14.6 sequence-search (36) to remove any that overlapped with previously annotated ncRNAs. Remaining novel microRNAs were added to the GTF file of known microRNAs, downloaded from miRBase v22 (35). Genome-mapped reads were assigned to mature microRNAs and quantified using featureCounts v1.6.0 (37) with the following settings: featureCounts -M -g gene_id -s 0.

### 2.3 RNA-seq analysis

BA2 and PBA/OFT E10.5 and E11.5 (25) and BA1 E10.5 and E11.5 RNA-seq libraries were adapter-trimmed and quality-filtered using Trimmomatic v0.36 (38). We used STAR v2.5.3a (39) to generate the mm10 genome index (GRCm38, release M23) and map RNA-seq reads to the genome using the mm10 genome primary fasta file and corresponding GTF file, both downloaded from GENCODE (40). Mapped reads were then assigned to annotated genomic features and quantified using featureCounts v1.6.0 (37), with options set as following: featureCounts -g gene_id -s 2.

### 2.4 Differential expression of small RNA and RNA-seq datasets

To remove lowly-expressed genes, a threshold was set for our RNA-seq libraries: 4.2CPM corresponding to approximately 100 reads for small RNA-seq, and 0.38CPM corresponding to approximately 10 reads for RNA-seq. Genes with CPM below these thresholds in two or more libraries were removed from further analysis. Differential analysis was performed using DESeq2 (41) and results were obtained for the following pairwise comparisons: E10.5 BA1 vs BA2, E10.5 BA1 vs PBA/OFT, E10.5 BA2 vs PBA/OFT, E11.5 BA1 vs BA2, E11.5 BA1 vs PBA/OFT, E11.5 BA2 vs PBA/OFT.

### 2.5 MicroRNA target prediction

3’UTR coordinates of BA transcripts were identified using the mm10 GTF and extract_transcript_regions.py (42). The longest 3’UTR for each gene was used, alongside BA expressed microRNAs, as inputs for *in silico* microRNA target prediction by seedVicious v1.3 (43). Results were filtered to remove predicted interactions with 6mers, off-6mers, sites with a hybridisation energy of >-7 kcal/mol, and 3’UTRs with <2 predicted microRNAs binding sites.

### 2.6 PBA/OFT gene set and microRNA-target enrichment analysis

E10.5 PBA/OFT enriched Gene Ontology (GO) terms were identified using gene set enrichment analysis by PANTHER (44). For input, we used 1.5-fold upregulated PBA/OFT genes (adj-p ≤0.05), with all pairwise BA 1.5-fold differentially expressed genes used as background. Genes that were annotated by ≥2-fold (FDR≤0.05) enriched GO terms were defined as our PBA/OFT gene set. To perform a hypergeometric test, we used the phyper function in R, with PBA/OFT subset interactions as the ‘sample’, and all BA interactions as the ‘population’. To identify the proportion of previously validated microRNA-target interactions, we used the multimiR package (45).

### 2.7 hESC cardiomyocyte differentiation

NKX2-5^eGFP/w^ hESCs (46) were seeded in hESC medium (DMEM/F-12 (Gibco, #31765027), 1X None-Essential Amino Acids (Gibco, #11140050), 1X GlutaMAX (Thermo Scientific, #35050038), 0.1mM 2-ME (Gibco, #21985023), 0.5% penicillin-streptomycin (Sigma-Aldrich, #P0781), 20% KnockOut Serum Replacement (KSR) (Gibco, 10828028), and 10ng/ml bFGF (Miltenyi, #130-104-924)) at a density of 1.8×10^5^ cells/ml on growth factor-reduced Matrigel coated 6-well plates. Twenty-four hours later (day 0) differentiation was induced as described previously (47). hESC medium was replaced with BPEL (48) supplemented with BMP4 (Bio-techne/R&D, #314-BP-050), 20ng/ml ACTIVIN A (Miltenyi Biotec, #130-115-009) and 1.75μM CHIR99021 (Selleckchem, #S1263). On day 3, media was refreshed with BPEL containing 1μM XAV939 (VWR, #CAYM13596-1). BPEL was refreshed every three days thereafter.

### 2.8 Cloning 3’UTRs into dual luciferase reporter vectors

Candidate target 3’UTRs were amplified using the following reaction; 1X Q5 Reaction Buffer (New England BioLabs, #B9027S), 20 units/ml Q5 High-fidelity DNA Polymerase (New England BioLabs, #M0491S), 0.6μM 3’UTR F/R primer (Supplementary Table 1), 0.2mM dNTP mix (Promega, #U1511), 2μl DNA, and RNase free water, using the cycling conditions: 98°C 1 minute, 35X [98°C 10 seconds, 58°C 30 seconds, 72°C 1 minute], 72°C 2 minutes. Amplified 3’UTRs were size selected and purified using the QIAquick Gel Extraction kit (Qiagen, #28704) following manufacturer’s instructions. Purified 3’UTRs were ligated with the pmirGLO dual-luciferase vector (Promega, #E1330) via SacI and SalI restriction sites. Resultant plasmids were sequenced using 0.4μM pmirGLO F/R custom sequencing primers (Supplementary Table 1).

### 2.9 MicroRNA 3’UTR binding site mutagenesis

miR-92b-3p binding sites in each 3’UTR were mutated using the QuikChange II XL site-directed mutagenesis kit (Agilent, #200517) following the manufacturer’s instructions, using custom mutagenesis primers (Supplementary Table 1). Mutated plasmids were sequenced as described above.

### 2.10 Dual luciferase reporter assay

NIH/3T3 cells were reverse co-transfected with 100ng pmirGLO dual luciferase vector (Promega, #E1330) containing appropriate 3’UTRs and 30nM microRNA mimic in 96-well plates (Invitrogen, miR-92b-3p #4464066, miRNA negative control 1 #4464058) using lipofectamine 2000 (Invitrogen, #11668019) diluted in opti-MEM (Gibco, #31985062). Luciferase activity was measured 48h later using the Dual-Glo Luciferase Assay System (Promega, #E2920) following the manufacturer’s instructions. Firefly and Renilla luciferase signal were measured using the Promega GloMax-Multi+ Detection System. Five technical replicates were performed for each biological replicate. Firefly luciferase values were first normalised to Renilla luciferase, and fold change was calculated relative to a control sample transfected with the dual-luciferase plasmid and no microRNA mimic.

### 2.11 MiR-92b-3p mimic transfection

HEK293 and NKX2-5^eGFP/w^ hESCs were reverse transfected with 30nM microRNA mimic (Invitrogen, miR-92b-3p #4464066, miRNA negative control 1 #4464058) using lipofectamine 2000 (Invitrogen, #11668019) diluted in opti-MEM (Gibco, #31985062). Samples were incubated for 24 hours at 37°C in 5% CO2.

### 2.12 RNA extraction and RT-qPCR

Where hESC cardiomyocyte differentiation timeline samples were used for microRNA and mRNA RT-qPCR, total RNA was extracted using the mirVana miRNA Isolation kit (Invitrogen, #AM1560) following the manufacturer’s instructions. Alternatively where only mRNA expression was measured, RNA was isolated using TRIzol Reagent (Invitrogen, #155996026) following a standard protocol. For miR-92b-3p quantification, 10ng total RNA was used as input with the TaqMan Advanced MiRNA cDNA Synthesis Kit (Applied Biosystems, #A28007) following the manufacturer’s instructions. For our internal control, U6, 10ng total RNA was used as input for the TaqMan MicroRNA Reverse Transcription Kit (Applied Biosystems, #4366596) following manufacturer’s instructions. To measure miR-92b-3p and U6 expression we used the TaqMan Fast Advanced Master Mix Kit (Applied Biosystems, #4444556) according to manufacturer’s instructions, with 1X TaqMan Advanced miRNA Assay (Applied Biosystems, #A25576, assay ID: 477823_mir) or 1X TaqMan small RNA Assay (Applied Biosystems, #4427975, assay ID: 001973) respectively. For mRNA expression we used the QuantiTect SYBR Green RT-PCR kit (Qiagen, #204243) in the following reaction: 1X QuantiTect SYBR Green RT-PCR master mix, 0.3μM F/R primers (Supplementary Table1), 1X RT mix, 40ng RNA, RNase-free water. Cycling conditions were: 50°C 30 minutes, 95°C 15 minutes, 40X [95°C 20 seconds, 57°C 30 seconds, 72°C 30 seconds], 68°C 7 minutes, 4°C hold. Fold change was calculated using 2^-ΔΔCt^ relative to the given control.

### 2.13 Western Blotting

HEK293 cells were lysed (20mM Tris-HCl, 120mM NaCl, 0.5mM EDTA, 0.5% NP40 (Thermo Scientific, #J60766-AP), 10% glycerol (Thermo Scientific, #17904), 1X protease cocktail inhibitor (Roche, #4693116001)) and the supernatant was recovered. Proteins were denatured in 1X laemmli buffer. Western blot membranes were incubated with 1:1500 anti-GATA-6 rabbit-mAb DEIE4 (Cellsignal, #5851) or 1:50,000 anti-β-Actin-peroxidase mouse-mAb (Sigma Aldrich, #A3854) in 1% milk. For GATA6 1:10,000 Goat anti-Rabbit IgG HRP (abcam, #ab6271) was used as a secondary antibody in 1% milk.

### 2.14 Sequence Alignment

miR-92 sequences for human, mouse, zebrafish and *Drosophila* were obtained from miRBase v22 (35). *Gata6* and *Tbx20* 3’UTR sequences were obtained from UCSC genome browser using hg38 and mm10, for human and mouse respectively. To align sequences we used Clustal Omega (49).

## 3. RESULTS

### 3.1 BAs demonstrate distinct global microRNA expression

To identify microRNAs that function during development and morphogenesis of mammalian BAs, we generated small-RNA-seq libraries for BA1, BA2, and PBA/OFT tissue from embryonic (E)10.5 and E11.5 mouse embryos. These datasets complement RNA-seq libraries we previously generated from equivalent tissues and timepoints (25), and so provide a relevant opportunity to consider expression of both microRNAs and their predicted target mRNAs.

Initial analysis demonstrated that BA small-RNA-seq libraries were enriched for 18-25nt reads (Supplementary Figure 1A), and 73-86% of the 18-25nt reads mapped to known microRNAs (Table 1). We additionally identified 28 novel pre-microRNAs (Supplementary Table 2), with two novel precursors each belonging to the mmu-miR-702 and mmu-miR-1839 families. As shown by principal component analysis (PCA) (Figure 1A), replicate samples clustered closely to one another indicating reproducibility. The first two principal components clearly separated the libraries based on their anterior-posterior location and developmental stage respectively. Using Spearman’s rank correlation, we found BA1 and BA2 E11.5 shared the greatest correlation, followed by PBA/OFT E10.5 and E11.5 (Supplementary Figure 1B). The PBA/OFT libraries were markedly distinct from the more anterior BA domains.

**Figure 1.**
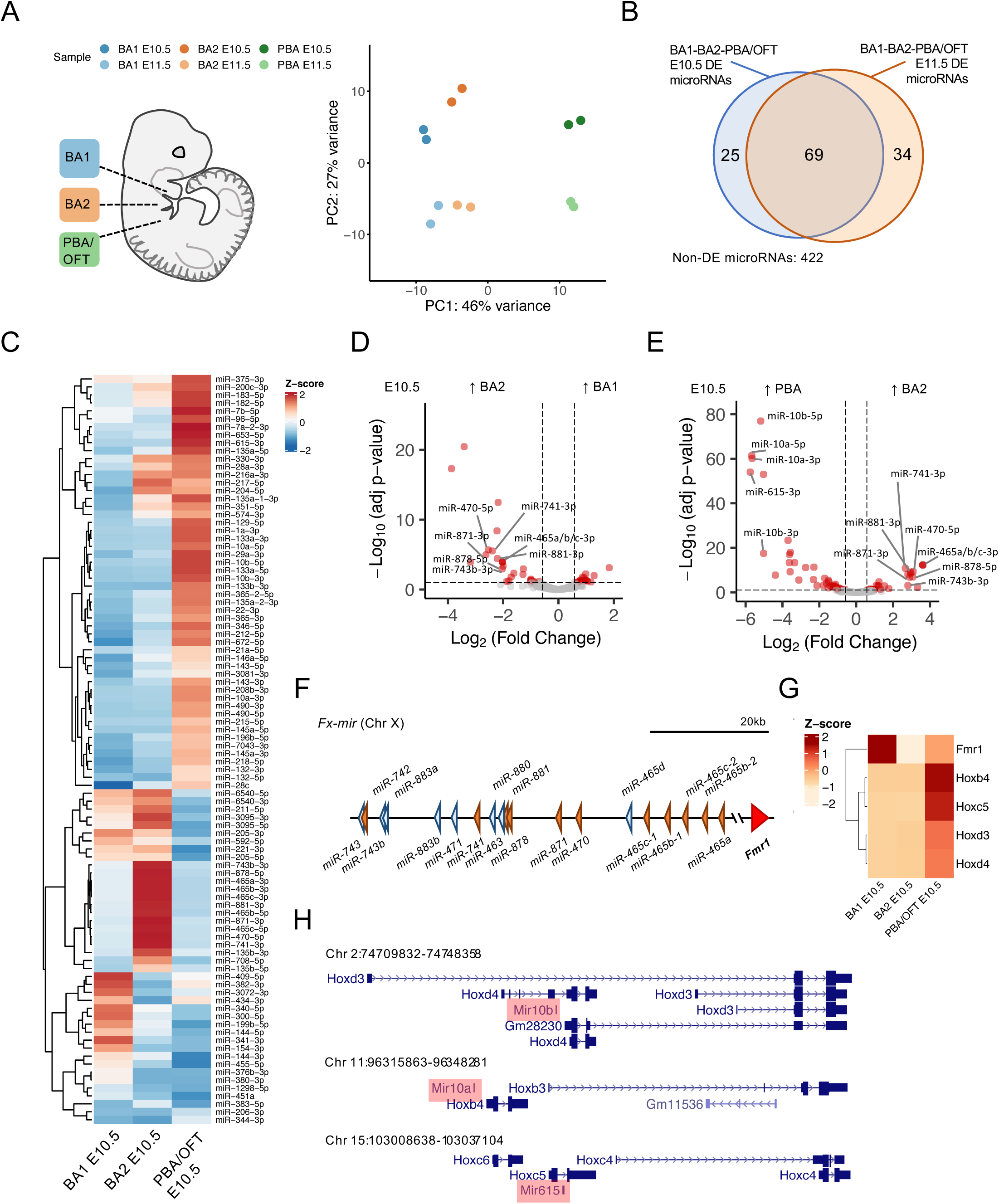
Differential expression of microRNAs across mouse BAs. A. Principal component analysis of branchial arch microRNA-seq libraries, mapped to the mm10 genome and assigned to mature microRNAs. B. Overlap of differentially expressed microRNAs (21.5-fold change and adj-p≤0.05), from BA pairwise comparisons at E10.5 and E11.5. C. Z-score normalised expression of differentially expressed microRNAs from E10.5 BA pairwise comparisons. D-E. Pairwise comparisons between E10.5 BA1 v BA2 and E10.5 BA2 v PBA/OFT. Fx-mir and Hox cluster microRNAs are labelled. Dotted lines correspond to 1.5-fold change and adjusted p-value≤0.05. F. *Fx-mir* cluster schematic. BA2 upregulated microRNAs are shaded orange and non-BA2 upregulated microRNAs are shaded blue. G. Gene expression of overlapping or nearby protein-coding genes from BA RNA-seq (25). H. Loci of PBA/OFT upregulated microRNAs overlapping Hox cluster gene members.

**Table 1.**
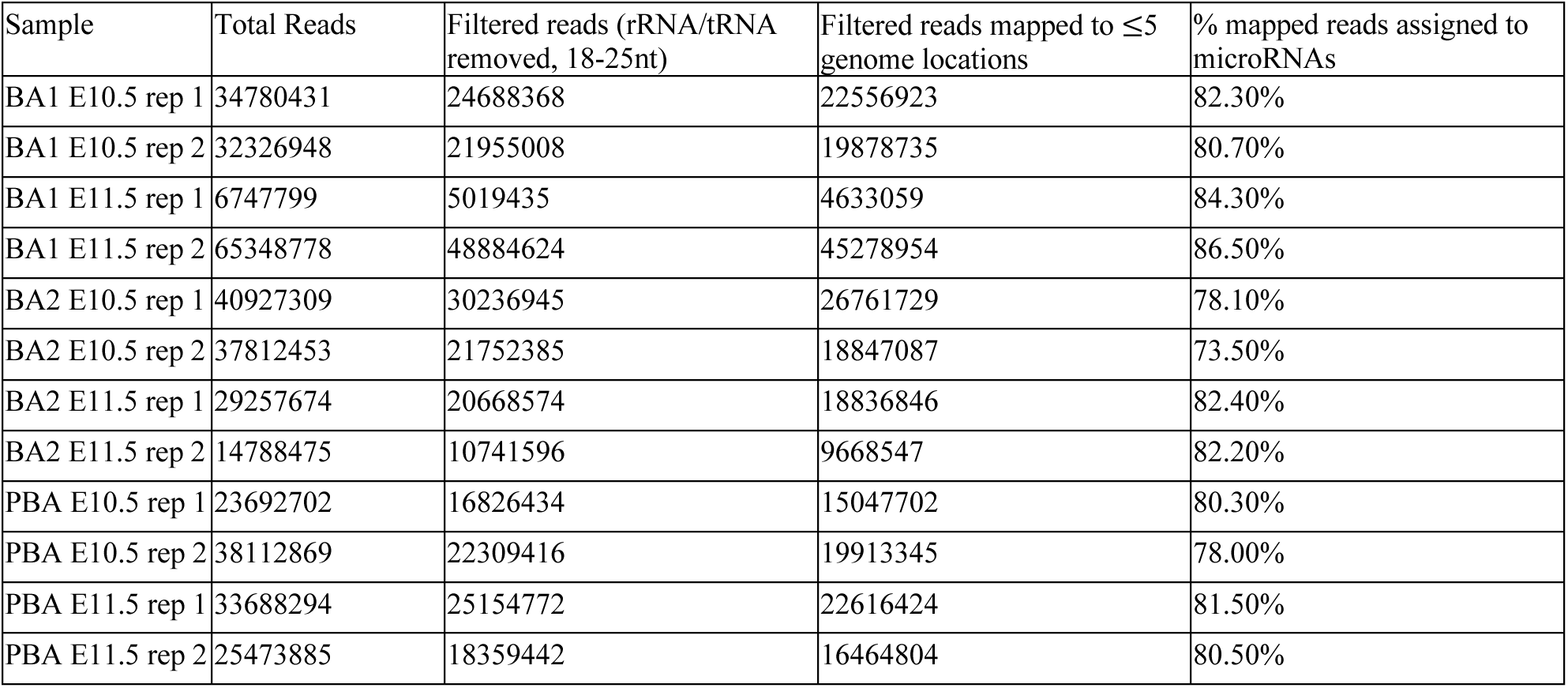
Mapping of small-RNA-sequencing libraries.

After applying an expression cut-off (see methods), we found a total of 550 microRNAs were expressed across the BAs. To identify tissue-specific microRNAs, we performed differential expression analysis on pairwise BA comparisons for E10.5 and E11.5 samples. We defined differentially expressed microRNAs as those with ≥1.5-fold change and adjusted p-value≤0.05. In at least one pairwise BA comparison for a given timepoint, 94 microRNAs were differentially expressed at E10.5, and 103 microRNAs were differentially expressed at E11.5. There was a large overlap between the sets of differentially expressed microRNAs at E10.5 and at E11.5 (Figure 1B), showing that BA-specific microRNA expression is maintained between these stages. Similarly, at both E10.5 and E11.5, the number of tissue-specific microRNAs increased across the anterior-posterior axis, with the majority of up-regulated microRNAs belonging to the PBA/OFT (Supplementary Figure 1C-D).

The combined expression of microRNAs and their target genes can provide an insight into microRNA function. For example, 11 mature microRNAs were significantly up-regulated in BA2 compared to both BA1 and PBA/OFT at E10.5 (Figure 1C). These are miR-743b-3p, miR-741-3p, miR-878-5p, miR-881-3p, miR-871-3p, miR-470-5p, miR-465a/b/c-3p, and miR-465b/c-5p (Figure 1D-E). All of these microRNAs are transcribed from a large microRNA cluster, *Fx-mir*, spanning ∼62kb on Chr X (Figure 1F). Members of this cluster have been described to regulate the neighbouring gene *Fmr1* (50,51), although each study draws contradictory conclusions with regards to whether these microRNAs repress or promote expression of *Fmr1*. Expression profiles from BA RNA-seq demonstrated that *Fmr1* was most lowly expressed in BA2 (Figure 1G), suggesting BA2 upregulated *Fx-mir* microRNAs may have a repressive role on *Fmr1* in BA2.

The PBA/OFT demonstrated the most distinct microRNA expression across the BAs, at both E10.5 and E11.5 (Supplementary Figure 1C-D). Some of the greatest upregulated microRNAs (Figure 1E, Supplementary Figure 1E) are located within the Hox clusters; *miR-10b* overlaps with both *Hoxd3* and *Hoxd4*, *miR-10a* is located upstream of *Hoxb4*, and *miR-615* is found within *Hoxc5* (Figure 1H). All of these *Hox* genes are also upregulated in the PBA/OFT E10.5 (Figure 1G), coinciding with spatial collinear *Hox* gene expression in distinct streams of cranial NC cells that populate the BAs (1). Therefore, Hox cluster microRNAs in the PBA/OFT mirror expression of their overlapping or nearby protein-coding genes.

The repertoire of tissue-specific microRNAs identified from BA pairwise comparisons were largely similar at E10.5 and E11.5 timepoints. As E10.5 samples demonstrated more defined PCA clustering (Figure 1A), and had a greater number of tissue-specific upregulated microRNAs (Supplementary Figure 1C-D), we focused on this timepoint for the remainder of our study.

### 3.2 miR-92b-3p is a candidate regulator of cardiac developmental genes

The microRNA biogenesis pathway has previously been implicated in PBA artery remodelling (21) and OFT morphogenesis (22). In addition, PBA/OFT microRNA expression is highly distinct compared to BA1 and BA2. Based on these findings, we set out to identify candidate PBA/OFT regulatory microRNAs, using the *in silico* microRNA-target prediction and gene set enrichment analysis approach shown in Figure 2A. To curate a subset of genes central to PBA/OFT development, we first identified PBA/OFT upregulated genes compared to both BA1 and BA2 (≥1.5-fold, adj-p≤0.05), giving us a total of 2078 genes. Using appropriate background genes (see methods), we performed gene set enrichment analysis on this subset to identify biological processes specific to PBA/OFT development. As shown in Figure 2B, GO terms that were enriched by ≥2-fold included many cardiac muscle development and differentiation related terms. This is consistent with the contribution of the PBA/OFT to cardiovascular development. Of the 2078 PBA/OFT upregulated genes, 315 were annotated under one or more of these enriched GO terms. We defined these 315 genes as our PBA/OFT cardiac subset.

**Figure 2.**
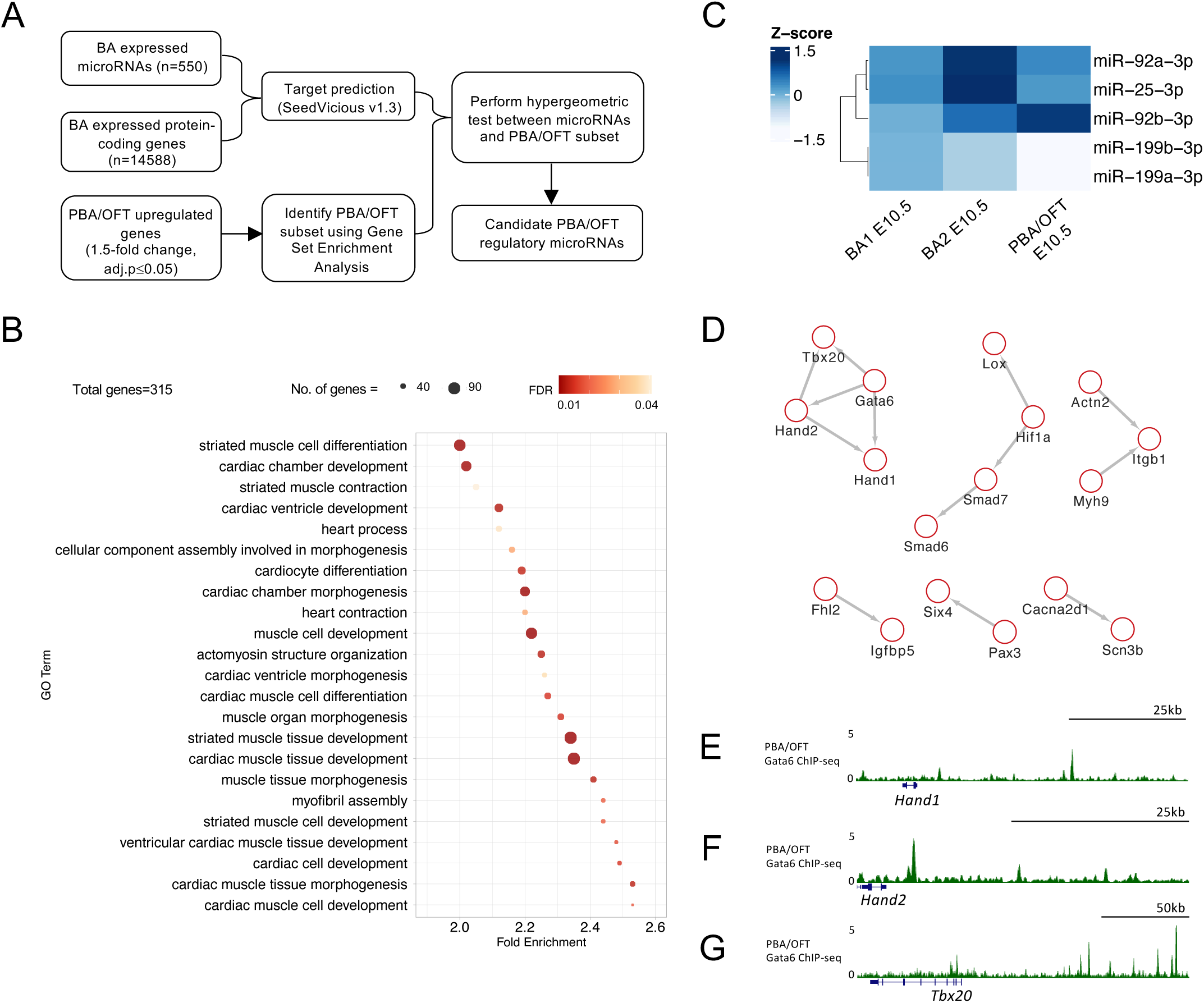
Identifying candidate microRNAs as regulators of PBA/OFT differentiation genes. A. Pipeline to identify candidate microRNA regulators of PBA/OFT differentiation. *In silico* microRNA-target predictions were generated for all BA expressed microRNAs and mRNAs. PBA/OFT upregulated genes were used to perform Gene Set Enrichment Analysis, to identify the PBA/OFT subset associated with enriched biological processes. Hypergeometric tests were performed to identify candidate regulatory microRNAs of the PBA/OFT subset. B. 2-fold enriched GO terms from Gene Set Enrichment Analysis. PBA/OFT upregulated genes from total RNA-seq were used as input, with all differentially expressed genes from BA pairwise comparisons used as background genes (fold-change21.5, adj-p≤0.05) (25). C. Normalized Z-score expression of the top five candidate microRNAs across the BAs at E10.5. D. Protein-protein interaction network of predicted miR-92b-3p targets included in the enriched GO terms in B, using STRING database with a 0.6 confidence interaction score. E-G. GATA6 binding sites around *Tbx20, Hand1,* and *Hand2* loci in mouse PBA/OFT (25).

Next, we identified microRNAs that were most significantly enriched for predicted interactions with our PBA/OFT cardiac subset. To achieve this, we performed hypergeometric tests using interactions from *in silico* microRNA-target predictions. The top five microRNAs, ranked by their hypergeometric p-value and predicted target enrichment were *miR-92b-3p*, *miR-199a-3p*, *miR199b-3p*, *miR-25-3p*, and *miR-92a-3p*. These microRNAs were 1.5-1.75-fold enriched for predicted interactions with our PBA/OFT subset, compared to predicted interactions with all BA expressed mRNAs. Additionally, a significantly higher percentage of these interactions were previously validated according to the multimiR database (45); 19% validated between the top-five ranked microRNAs and the PBA/OFT gene subset, compared to 5% for all BA microRNAs and the PBA/OFT subset.

Next, we considered the expression of the top-five ranked microRNAs across the BAs (Figure 2C). The highest ranked microRNA, miR-92b-3p, demonstrated higher expression in the PBA/OFT. In contrast, the other four microRNAs either displayed higher expression in BA2: miR-92a-3p and miR-25-3p, or BA1: miR-199a/b-3p. MicroRNAs can be either positively correlated or inversely correlated in expression with their targets, depending on the network motif they function within (52,53). Interestingly, miR-25-3p and miR-92a/b-3p share the same seed sequence, suggesting this microRNA family could collectively regulate the PBA/OFT cardiac subset, although potentially via alternative network motifs. We focused on miR-92b-3p, the top ranked microRNA predicted to target the PBA/OFT subset and upregulated in this domain. miR-92b-3p was predicted to bind to 30 of the 315 genes in the PBA/OFT subset (Supplementary Table 3). We visualised protein-protein interactions to identify functional connectivity between miR-92b-3p predicted targets and found multiple clusters of proteins (Figure 2D). Four cardiac TFs; GATA6, TBX20, HAND1 and HAND2, clustered together, and share reported direct and indirect functional links (54). In *GATA6* loss of function human induced pluripotent stem cells (hiPSCs) undergoing cardiomyocyte differentiation, *HAND1, HAND2* and *TBX20* were downregulated (54). Supporting direct regulation of *HAND1*, *HAND2* and *TBX20* by GATA6, we found prominent GATA6 peaks within 100kb upstream or downstream of *Hand1*, *Hand2*, and *Tbx20* in the PBA/OFT (25) (Figure 2E-G). Taken together, these four TFs may belong to a cardiac GRN. MicroRNAs are understood to stabilise GRNs through broad regulation of multiple targets (55). As *Hand2* is a validated target of miR-92b-3p (26), we next explored miR-92-3p regulation of the other predicted targets *Gata6*, *Tbx20* and *Hand1*.

### 3.3 miR-92b-3p interacts with *Gata6* and *Tbx20* 3’UTRs

We assessed whether miR-92b-3p can interact with its predicted target sites in *Gata6*, *Tbx20* and *Hand1* 3’UTRs, using dual-luciferase reporter assays. Portions of each 3’UTR containing the predicted binding sites were cloned into the Promega pmirGLO dual-luciferase plasmid (Figure 3A). Following co-transfection of miR-92b-3p mimic and *Gata6* and *Tbx20* 3’UTR reporters, we saw a significant reduction in luciferase signal, approximately 0.5-fold, compared to the negative control mimic (Figure 3B-C). In contrast, when we co-transfected miR-92b-3p with the *Hand1* 3’UTR reporter, we saw a much smaller reduction in reporter signal compared to the negative control (Figure 3D). These results are consistent with the predicted hybridisation energies between miR-92b-3p and the three target sites (Figure 3E-G), whereby the *Hand1* 3’UTR target site is the weakest predicted at -7.2kcal/mol, just meeting our -7kcal/mol threshold cut-off.

**Figure 3.**
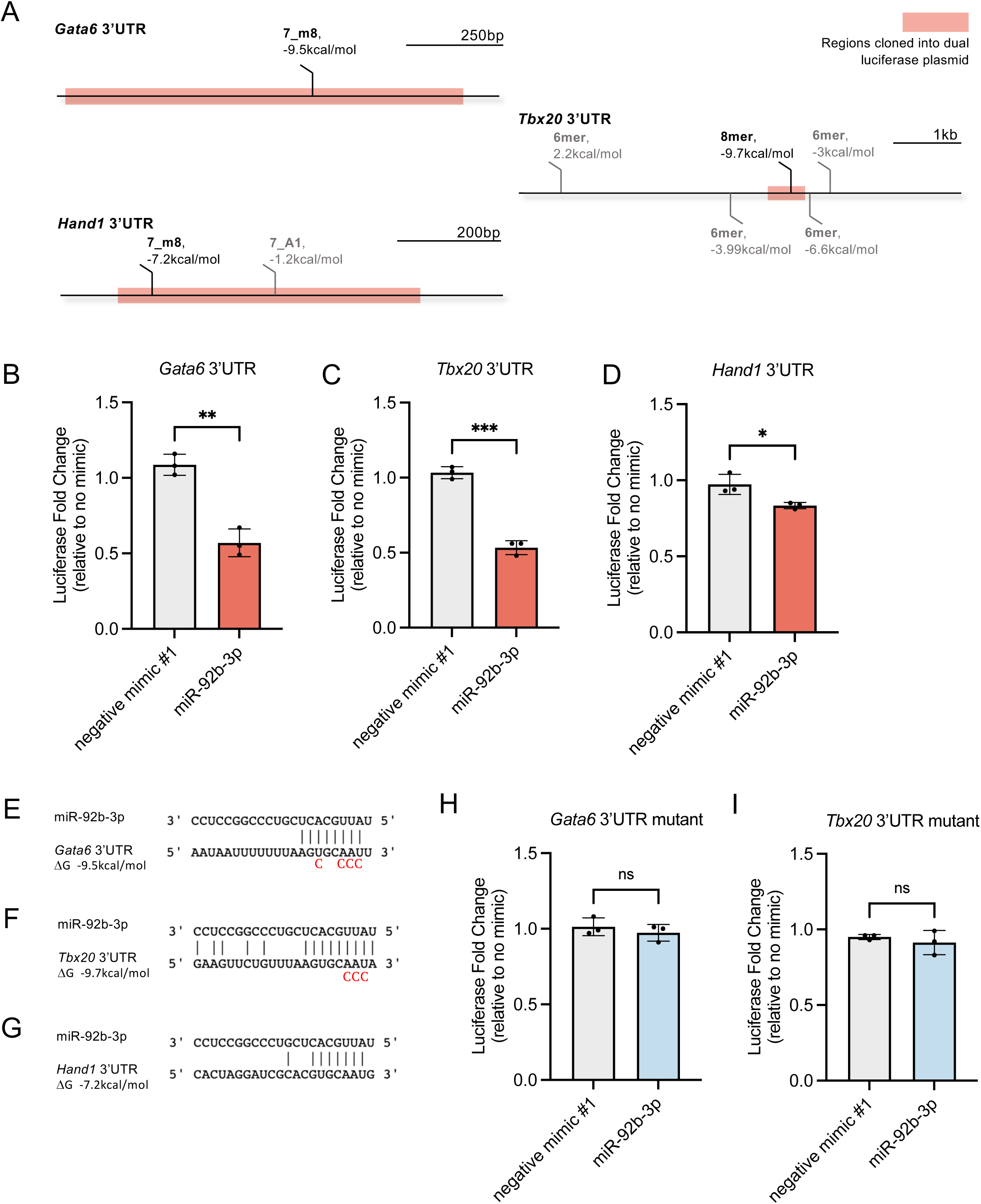
Testing miR-92b-3p binding sites in mouse *Gata6, Tbx20,* and *Hand1* 3’UTRs. A. Seedvicious predicted binding sites between miR-92b-3p and *Gata6, Tbx20*, and *Hand1* 3’UTR’s. Highlighted regions were cloned into the pmirGLO dual-luciferase plasmid. Sites that did not meet our threshold of -7kcal/mol are shown in grey. B-D. Dual-luciferase reporter assays following co-transfection of microRNA mimics and dual luciferase plasmids containing wild type 3’UTRs. Luciferase fold change is plotted relative to samples transfected with only plasmid and no mimic. Negative mimic #1 is ‘mirVana negative control #1’. Values are presented as the mean±s.d, n=3 biological replicates. Significance was calculated using an unpaired t-test: *Gata6* p-value=0.0015, *Tbx20* p-value=0.001, *Hand1* p-value=0.0254. E-G. Predicted complementary binding and hybridisation energy between miR-92b-3p and *Gata6, Tbx20, Hand1* 3’UTRs. Mutated nucleotides included underneath wildtype sequences. H-I. Dual-luciferase reporter assays following co-transfection of dual luciferase plasmids containing mutated 3’UTRs microRNA mimics. Values are presented as mean±s.d., n=3 biological replicates.

Next, we determined whether the luciferase reporter knockdown was specifically caused by the physical interaction of miR-92b-3p with its putative binding sites in *Gata6* and *Tbx20* 3’UTRs. We mutated the nucleotides complementary to miR-92b-3p’s seed region (Figure 3E-F) to disrupt initial binding, which reportedly occurs between a microRNA’s seed site, at positions 2-4, and its target RNA before extended base pairing 3’ of this region (56). Following co-transfection of mutated 3’UTRs with miR-92b-3p, we did not observe any reduction in luciferase signal, confirming luciferase signal knockdown requires interaction of miR-92b-3p with its binding sites in *Gata6* and *Tbx20* 3’UTRs (Figure 3H-I). In summary, miR-92b-3p can specifically interact with the predicted targets, and the extent of repression is reflective of the predicted hybridisation energy.

### 3.4 miR-92b-3p binding to *Gata6* and *Tbx20* is conserved in human

miR-92b-3p is conserved between species separated by more than 780 million years of evolution (57). Using Clustal Omega (49), alignment of *pre-miR-92b* across mouse, human, zebrafish, and *Drosophila* showed the majority of conserved nucleotides are located in the 3’ mature arm (Figure 4A). Importantly, the validated miR-92b-3p binding sites in mouse *Gata6* and *Tbx20* 3’UTRs are located in regions of high sequence conservation (58) (Figure 4B-C, Supplementary Figure 2A-B). Consistently, *in silico* target prediction between human miR-92b-3p and *GATA6* and *TBX20* 3’UTRs identified homologous binding sites to those we identified by luciferase reporter assay, with identical seed complementarity and similar hybridisation energies (Supplementary Figure 2C-D). Therefore, we wanted to determine whether miR-92b-3p binding to *GATA6* and *TBX20* was conserved to human.

**Figure 4.**
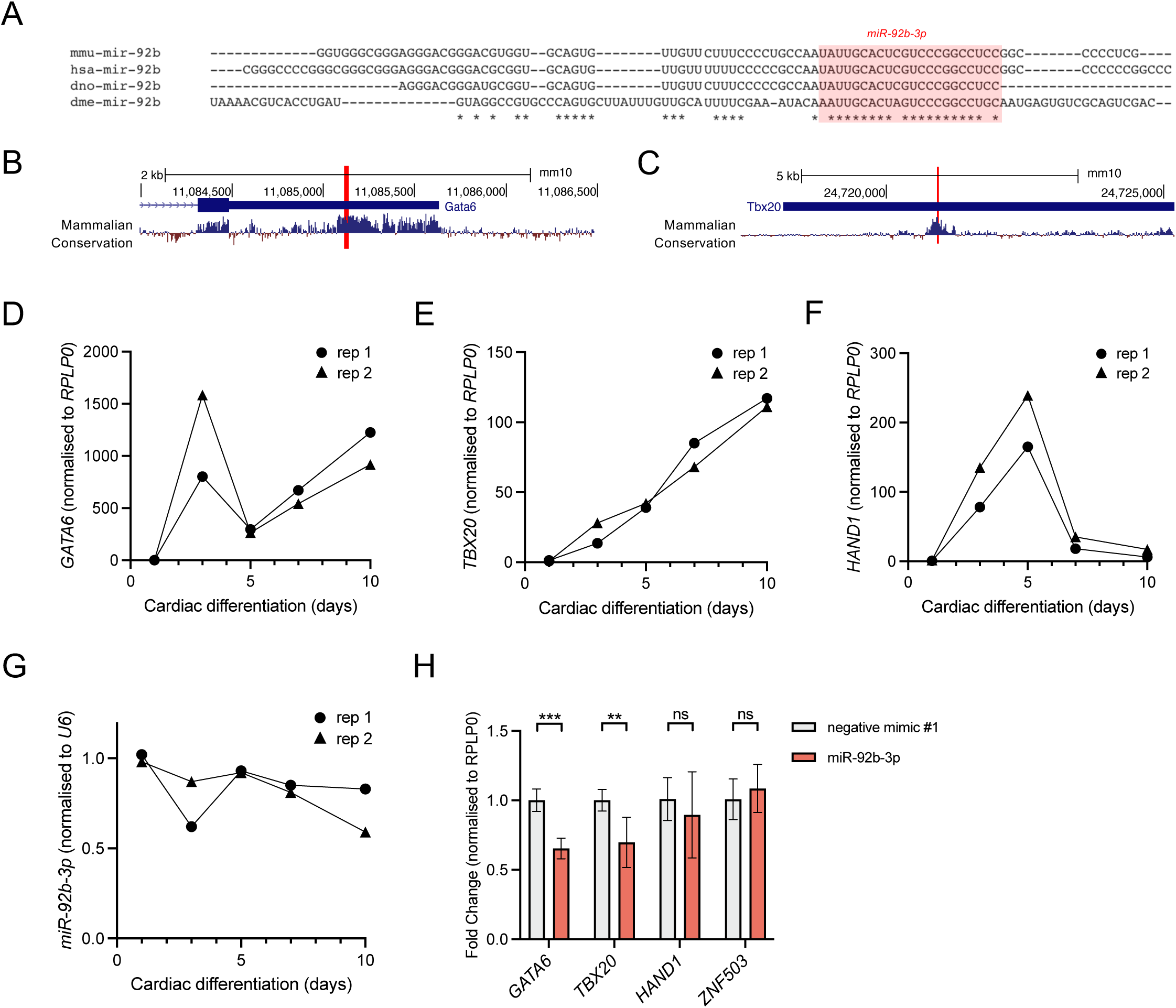
Conservation of miR-92b-3p targeting of *GATA6* and *TBX20*. A. Alignment of *miR-92b* (3p highlighted) in *M. musculus*, *H. sapiens*, *D. rerio*, and *D. melanogaster*. B-C. PhyloP placental mammalian basewise conservation at *Gata6* and *Tbx20* 3’UTRs. miR-92b-3p binding sites are highlighted in each 3’UTR. D-F. RT-qPCR of *GATA6*, *TBX20* and *HAND1* expression, normalised to *RPLP0*, during days 1-10 of hESC cardiomyocyte differentiation, n=2 biological replicates. (H) *miR-92b-3p* expression, normalised to snRNA *U6*, during days 1-10 of a hESC cardiomyocyte differentiation, n=2 biological replicates. (I) RT-qPCR of *GATA6*, *TBX20, HAND1,* and *ZNF503* (negative control), following 24h transfection with miR-92b-3p mimic. hESCs were collected on day 7. Expression was normalised to *RPLP0* and then used to calculate fold change relative to negative mimic #1 control. Values are presented as the mean±s.d., n=6 biological replicates. Statistical significance was calculated performing multiple unpaired t-tests, *GATA6* adjusted p-value = 8.4×10^-5^, *TBX20* adjusted p-value = 4.6×10^-3^.

As the PBA/OFT transcriptome is enriched for transcripts linked to cardiac muscle development and differentiation (Figure 2B), we employed a human cardiac differentiation system (47). Firstly, we determined the expression dynamics of both miR-92b-3p and its identified targets *GATA6* and *TBX20*, in addition to *HAND1*. All three TFs were initially upregulated at the onset of differentiation, however, from day 3 onwards they displayed variable expression dynamics (Figure 4D-F). *GATA6* and *TBX20* expression increased between day 5 to day 10, whilst *HAND1* expression decreased. In contrast, we found miR-92b-3p remained generally stable throughout cardiac differentiation, and slightly decreased by day 10 (Figure 4G). We also measured miR-92b-3p in undifferentiated hESCs and found it to be expressed at similar levels as day 1 (data not shown), demonstrating that its expression was not induced by differentiation. This is comparable to microarray data previously published (59). Therefore, whilst not expressed in a cardiac specific manner, miR-92b-3p is expressed at the same time as its identified targets *GATA6* and *TBX20*, and could therefore function as a regulatory factor in a cardiac setting.

To test whether miR-92b-3p can regulate endogenous *GATA6* and *TBX20,* we transfected differentiating cardiomyocyte cells with microRNA mimics and measured gene expression 24 hours later. *GATA6* and *TBX20* were both downregulated in the miR-92b-3p transfected samples compared to the negative mimic control (Figure 4H), indicating miR-92b-3p can downregulate their expression in a cardiac setting. In contrast, we found no significant knockdown of *HAND1.* Additionally, a negative control gene *ZNF503* (with no predicted miR-92b-3p binding sites in its 3’UTR) did not display significant knockdown between conditions, demonstrating that miR-92b-3p transfection did not affect global transcription or translational machinery.

As *GATA6* loss of function has previously been shown to downregulate *TBX20* in a cardiac setting (54), the *TBX20* knockdown we observe could be an indirect result, rather than a direct effect of miR-92b-3p. However, the luciferase reporter assays do indicate miR-92b-3p interacts with the *Tbx20* 3’UTR. To distinguish between these possibilities, we probed GATA6 and *TBX20* functional relationship in a non-cardiogenic context. We overexpressed GATA6 in HEK293 cells and measured gene expression 48h later. *GATA6* was approximately 1000-fold higher, with no significant change in *TBX20* (Supplementary Figure 2E), suggesting that GATA6 does not regulate *TBX20* expression in this cell type. Therefore, we measured endogenous *GATA6* and *TBX20* in HEK293 cells following miR-92b-3p transfection (Supplementary Figure 2F). Both *GATA6* and *TBX20* were significantly knocked down following miR-92b-3p transfection. Furthermore, we found reduced GATA6 expression, showing that miR-92b-3p repression extended to protein abundance (Supplementary Figure 2G). We did not see any significant difference in our negative control, ACTB, for both mRNA and protein, again showing that overexpression of miR-92b-3p is unlikely to disrupt global transcription and translation. Combined with the ability of miR-92b-3p to specifically target *Gata6* and *Tbx20* 3’UTRs, these results indicate that miR-92b-3p can independently target both *GATA6* and *TBX20* to regulate their expression. This places miR-92b-3p within a cardiac GRN, whereby it can regulate the network at multiple levels, likely providing stability and promoting developmental canalisation.

## 4. DISCUSSION

In this study we present novel microRNA datasets characterising global expression across developing mammalian BAs as they undergo tissue specification and morphological changes. These libraries complement our previously published work on RNA-seq and ChIP-seq datasets (25,60,61), expanding our understanding of BA developmental biology into the microRNA field. These microRNA-seq datasets provide information on microRNA expression across all BA domains, building on microRNA microarray data generated from isolated NC cells in mouse BA1 (62). None of the NC-upregulated microRNAs Sheehy et al., (2010) reported on demonstrated BA1-specific expression in our datasets, suggesting alternative groups of microRNAs may regulate whole BA identity compared to distinct cell populations within the BAs.

We have characterised expression of 550 mature microRNAs in the BAs, with the most distinct domain being the PBA/OFT with regards to microRNA upregulation. Additionally, we identified miR-92b-3p as a candidate regulator of cardiovascular development in the PBA/OFT. We validated its interaction with *Gata6* and *Tbx20,* two central cardiac TFs (25,54,63,64). Previously, knockout of the microRNA biogenesis factor *Dicer* led to abnormal OFT development, with progenitor cells failing to differentiate into smooth muscle cells (SMC) (62). Incidentally, Gata6 is sufficient to promote SMC differentiation (25), highlighting one of the many microRNA-target interactions that may support normal OFT development. Cardiac development is also understood to be sensitive to gene or protein dosage, echoed by the incidence of human congenital cardiac malformations (65). Therefore, it is understandable that cardiovascular development is in part controlled by microRNA-directed regulation, and that microRNA dysregulation can therefore have an impact on disease (27,66–68).

TFs have been described as microRNA target hubs, with microRNAs essentially “regulating the regulators” (69). Previously the cardiac TF *Hand2* was identified as a target of miR-92b-3p (26), consistent with our *in silico* target predictions. Both Yu et al., (2019) and Hu et al., (2017) showed that Ang-II induced cardiomyocyte hypertrophy caused an increase in miR-92b-3p expression in neonatal mouse ventricular cells. Furthermore, overexpression of miR-92b-3p prevented the hypertrophic phenotype developing following Ang-II treatment, through targeting *Hand2* (26). Another cardiac TF, *Mef2d*, is targeted by miR-92b-3p (27,28). While we identified *Mef2d* in our predicted targets of miR-92b-3p, *Mef2d* was not annotated under any of the GO terms we found enriched for PBA/OFT upregulated genes and therefore not included in our PBA/OFT gene set.

The understanding of microRNA regulation within GRNs has advanced since the application of computational and mathematical modelling approaches (53,70). Different GRN motifs elicit different outputs (53), and it is therefore important to consider where a microRNA fits into a GRN to infer its functional role. GATA6, TBX20 and HAND2 are all cardiac progenitor markers and have important roles in activating cardiovascular cell fates (25,54,63,64,71). Furthermore, these three TFs demonstrate overlapping expression within the PBA/OFT, as shown by in situ hybridisation and fluorescence microscopy (25,64,72,73). GATA6 indirectly promotes *HAND2* and directly promotes *TBX20* expression during hiPSC cardiomyocyte differentiation, through functioning as a pioneer cardiac factor (54). Additionally, HAND2 binds to cis-regulatory modules associated with *Gata6* and *Tbx20* in embryonic hearts (74). Taken together, this places miR-92b-3p in multiple microRNA-mediated coherent feedforward loops (Figure 5). Of note, the miR-92b-3p target sites in *Gata6*, *Tbx20* and *Hand2* 3’UTRs are all located in highly conserved regions, determined from the UCSC genome browser Multiz alignments of 60 vertebrates, suggesting these are fundamental networks. These networks can function to minimise leaky transcripts or prevent spatial co-expression of the microRNA and its targets (53,75). From our microRNA-seq and RNA-seq datasets we know that miR-92b-3p, *Gata6* and *Tbx20* are all upregulated in the PBA/OFT domain. However, it would be interesting to determine at a greater resolution, for example single-cell, whether miR-92b-3p is in fact inversely correlated with *Gata6, Tbx20* and *Hand2,* as may be expected if functioning to prevent spatial co-expression or minimise leaky transcripts (53).

**Figure 5.**
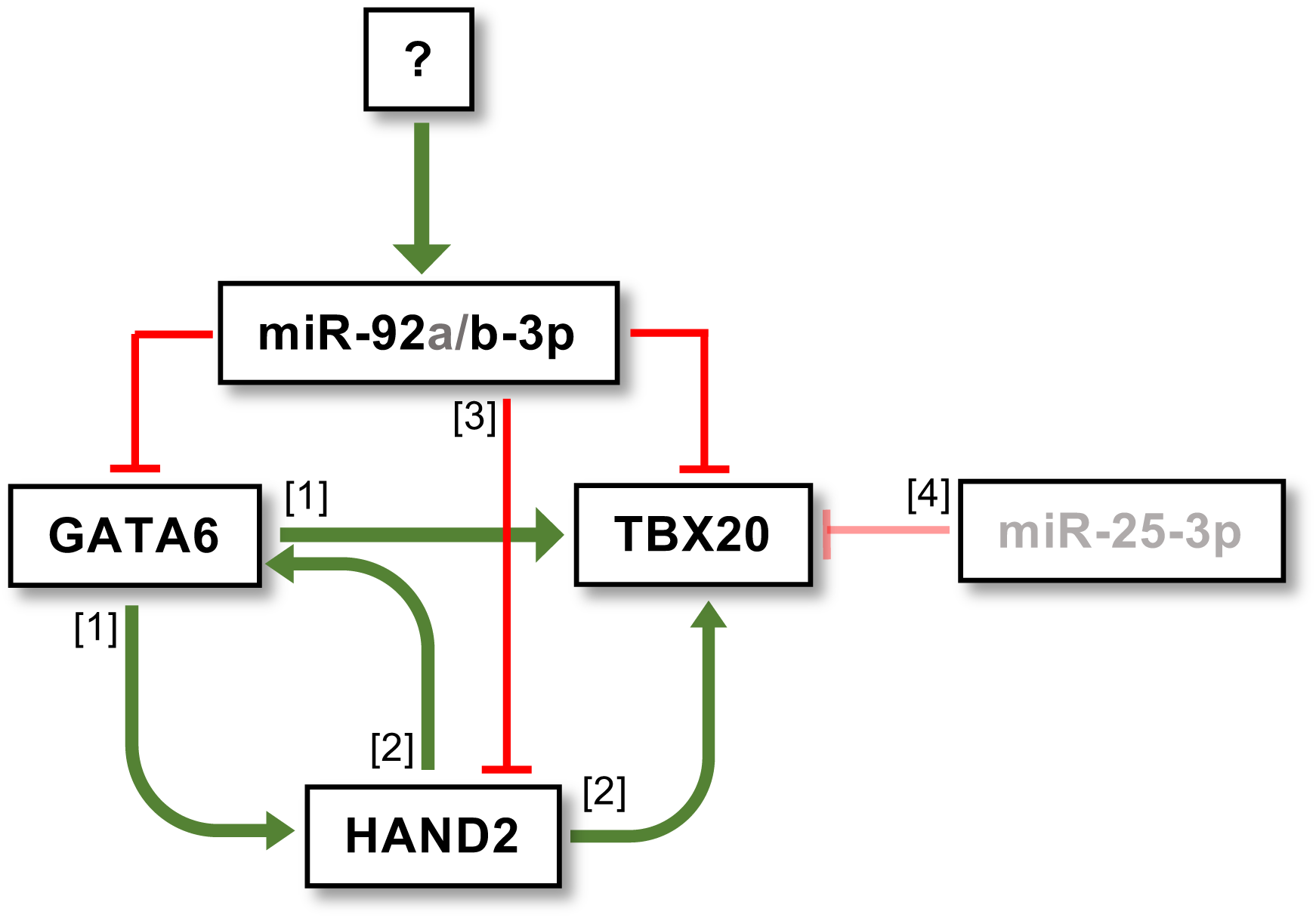
MicroRNA-mediated coherent feedforward loops which miR-92b-3p could regulate during PBA/OFT development. Regulatory interactions between TFs and microRNAs are supported by the following studies: [1] Sharma et al., 2020 [2] Laurent et al., 2017, [3] Yu et al., 2019, [4] Alzein et al., 2021.

Alternatively, microRNAs can also work as master regulators when embedded within coherent feedforward motifs (70). If the microRNA and target genes are expressed in the same cell, then the microRNA concentration is a controlling parameter, driven by competition for microRNA-target binding. The microRNA in turn can regulate and maintain the ratio of its targets relative to one another, ensuring stability in target concentration (70). This is particularly effective when one of the microRNA targets is a TF which regulates the other target (76), as we see in our miR-92b-3p network, where GATA6 regulates *Tbx20.* We have evidence of *Gata6* and *Tbx20* co-expression in human embryonic and fetal OFT single-cell data (unpublished). Therefore, if miR-92b-3p were also co-expressed in these cells, microRNA-target competition could occur. As a result, miR-92b-3p could act to reinforce this GRN and facilitate PBA/OFT development in a “coordinate regulatory” manner (55). To build on our understanding of miR-92b-3p within this GRN, it would be interesting to determine its upstream regulator.

MicroRNAs often act moderately to fine-tune their target gene expression, with the idea that “weak and broad” regulation is central to how microRNAs stabilise GRNs and contribute to developmental canalization (55,77). However, this moderately repressive role often means there is no substantial phenotypic consequence when individual microRNAs are knocked out, as over 90% of microRNA activity is recognised as “weak” (18). There is also redundancy between microRNAs that share the same targets, therefore it is also important to consider how microRNAs may work collectively. miR-92b-3p belongs to a larger seed family of microRNAs containing miR-92a-1/2-3p, miR-25-3p, and miR-363-3p. Two of these, miR-92a-3p and miR-25-3p, were also in the top five candidates predicted to regulate our PBA/OFT gene subset in this study. As these microRNAs share identical seed sequences, there will likely be a high level of redundancy between their targets (78,79). miR-92a-3p is located within the *miR-17-92* cluster, which has previously been linked to cardiomyocyte proliferation, hypertrophic cardiomyopathy, and aberrant cardiac ageing (80–82). Furthermore, deletion of this cluster caused ventricular septal defects in mouse models (66). Another family member, miR-25, is expressed in the OFT and ventricular regions during embryonic chick development, and predicted to regulate *Tbx20* (14). Evidence for miR-92b-3p regulated cardiac development extends to *Drosophila*, whereby miR-92b-3p exhibited muscle and cardiac specific expression (28). Taken together, we hypothesise that miR-92b-3p and members of its family, through cooperativity and redundancy, perform a central role in regulating the described cardiac GRN (Figure 5) during PBA/OFT development.

## DATA AVAILABILITY

Fastq files corresponding to small RNA-sequencing libraries are available from the European Nucleotide Archive: ERR12176799 BA1_E10.5_Rep1, ERR12176800 BA2_E10.5_Rep1, ERR12176801 PBA_E10.5_Rep1, ERR12176802 BA1_E10.5_Rep2, ERR12176803 BA2_E10.5_Rep2, ERR12176804 PBA_E10.5_Rep2, ERR12176805 BA1_E11.5_Rep1, ERR12176806 BA2_E11.5_Rep1, ERR12176807 PBA_E11.5_Rep1, ERR12176808 BA1_E11.5_Rep2, ERR12176809 BA2_E11.5_Rep2, ERR12176810 PBA_E11.5_Rep2.

## SUPPLEMENTARY DATA

Supplementary data is included with this submission.

## ACKNOWLEDGEMENTS

We would like to acknowledge the Genomic Core Technology Facility at The University of Manchester for performing the small-RNA sequencing. We would also like to thank Joshua Mallen for advice and technical troubleshooting, Matthew Birket for advice on hESC cardiomyocyte differentiation, and Emma Layton and Svitlana Kurinna for help with microRNA assays.

## FUNDING

This work was supported by Wellcome Trust [R121377].

## CONFLICT OF INTEREST

The authors declare they have no conflict of interest.

## Notes

https://www.ebi.ac.uk/ena/browser/view/PRJEB64007

### Competing Interest Statement

The authors have declared no competing interest.

## REFERENCES

1. Frisdal, A. and Trainor, P.A. (2014) Development and evolution of the pharyngeal apparatus. Wiley interdisciplinary reviews. Developmental biology, 3, 403–418.

2. Graham, A. and Richardson, J. (2012) Developmental and evolutionary origins of the pharyngeal apparatus. EvoDevo, 3, 24.

3. Kulesa, P.M., Bailey, C.M., Kasemeier-Kulesa, J.C. and McLennan, R. (2010) Cranial neural crest migration: new rules for an old road. Dev Biol, 344, 543–554.

4. Etchevers, H.C., Dupin, E. and Le Douarin, N.M. (2019) The diverse neural crest: from embryology to human pathology. Development, 146.

5. Kirby, M.L., Turnage III, K.L. and Hays, B.M. (1985) Characterization of conotruncal malformations following ablation of “cardiac” neural crest. The Anatomical Record, 213, 87–93.

6. Boot, M.J., Gittenberger-De Groot, A.C., Van Iperen, L., Hierck, B.P. and Poelmann, R.E. (2003) Spatiotemporally separated cardiac neural crest subpopulations that target the outflow tract septum and pharyngeal arch arteries. The Anatomical Record Part A: Discoveries in Molecular, Cellular, and Evolutionary Biology, 275, 1009–1018.

7. Kirby, M.L., Gale, T.F. and Stewart, D.E. (1983) Neural crest cells contribute to normal aorticopulmonary septation. Science, 220, 1059–1061.

8. Bergwerff, M., Verberne, M.E., DeRuiter, M.C., Poelmann, R.E. and Gittenberger-de-Groot, A.C. (1998) Neural Crest Cell Contribution to the Developing Circulatory System. Circulation Research, 82, 221–231.

9. Hiruma, T., Nakajima, Y. and Nakamura, H. (2002) Development of pharyngeal arch arteries in early mouse embryo. J Anat, 201, 15–29.

10. Jiang, X., Rowitch, D.H., Soriano, P., McMahon, A.P. and Sucov, H.M. (2000) Fate of the mammalian cardiac neural crest. Development, 127, 1607–1616.

11. Gurjarpadhye, A., Hewett, K.W., Justus, C., Wen, X., Stadt, H., Kirby, M.L., Sedmera, D. and Gourdie, R.G. (2007) Cardiac neural crest ablation inhibits compaction and electrical function of conduction system bundles. Am J Physiol Heart Circ Physiol, 292, H1291–1300.

12. Soldatov, R., Kaucka, M., Kastriti, M.E., Petersen, J., Chontorotzea, T., Englmaier, L., Akkuratova, N., Yang, Y., Häring, M., Dyachuk, V. et al. (2019) Spatiotemporal structure of cell fate decisions in murine neural crest. Science, 364, eaas9536.

13. Li, C.-J., Liau, E.S., Lee, Y.-H., Huang, Y.-Z., Liu, Z., Willems, A., Garside, V., McGlinn, E., Chen, J.-A. and Hong, T. (2021) MicroRNA governs bistable cell differentiation and lineage segregation via a noncanonical feedback. Molecular Systems Biology, 17, e9945.

14. Alzein, M., Lozano-Velasco, E., Hernández-Torres, F., García-Padilla, C., Domínguez, J.N., Aránega, A. and Franco, D. (2021) Differential Spatio-Temporal Regulation of T-Box Gene Expression by microRNAs during Cardiac Development. J Cardiovasc Dev Dis, 8.

15. Crist, C.G., Montarras, D., Pallafacchina, G., Rocancourt, D., Cumano, A., Conway, S.J. and Buckingham, M. (2009) Muscle stem cell behavior is modified by microRNA-27 regulation of Pax3 expression. Proceedings of the National Academy of Sciences, 106, 13383–13387.

16. Aboobaker, A.A., Tomancak, P., Patel, N., Rubin, G.M. and Lai, E.C. (2005) *Drosophila* microRNAs exhibit diverse spatial expression patterns during embryonic development. Proceedings of the National Academy of Sciences, 102, 18017–18022.

17. Avital, G., França, G.S. and Yanai, I. (2017) Bimodal Evolutionary Developmental miRNA Program in Animal Embryogenesis. Molecular Biology and Evolution, 35, 646–654.

18. Chen, Y., Shen, Y., Lin, P., Tong, D., Zhao, Y., Allesina, S., Shen, X. and Wu, C.I. (2019) Gene regulatory network stabilized by pervasive weak repressions: microRNA functions revealed by the May-Wigner theory. Natl Sci Rev, 6, 1176–1188.

19. Ma, F., Lin, P., Chen, Q., Lu, X., Zhang, Y.E. and Wu, C.-I. (2018) Direct measurement of pervasive weak repression by microRNAs and their role at the network level. BMC genomics, 19, 1–12.

20. Zhao, Y., Lin, P., Liufu, Z., Yang, H., Lyu, Y., Shen, X., Wu, C.-I. and Tang, T. (2018) Regulation of Large Number of Weak Targets—New Insights from Twin-microRNAs. Genome biology and evolution, 10, 1255–1264.

21. Nie, X., Wang, Q. and Jiao, K. (2011) Dicer activity in neural crest cells is essential for craniofacial organogenesis and pharyngeal arch artery morphogenesis. Mech Dev, 128, 200–207.

22. Saxena, A. and Tabin, C.J. (2010) miRNA-processing enzyme Dicer is necessary for cardiac outflow tract alignment and chamber septation. Proc Natl Acad Sci U S A, 107, 87–91.

23. Sellier, C., Hwang, V.J., Dandekar, R., Durbin-Johnson, B., Charlet-Berguerand, N., Ander, B.P., Sharp, F.R., Angkustsiri, K., Simon, T.J. and Tassone, F. (2014) Decreased DGCR8 Expression and miRNA Dysregulation in Individuals with 22q11.2 Deletion Syndrome. PLOS ONE, 9, e103884.

24. Gregory, R.I., Yan, K.-p., Amuthan, G., Chendrimada, T., Doratotaj, B., Cooch, N. and Shiekhattar, R. (2004) The Microprocessor complex mediates the genesis of microRNAs. Nature, 432, 235–240.

25. Losa, M., Latorre, V., Andrabi, M., Ladam, F., Sagerström, C., Novoa, A., Zarrineh, P., Bridoux, L., Hanley, N.A., Mallo, M., et al. (2017) A tissue-specific, Gata6-driven transcriptional program instructs remodeling of the mature arterial tree. eLife, 6, e31362.

26. Yu, X.-J., Huang, Y.-Q., Shan, Z.-X., Zhu, J.-N., Hu, Z.-Q., Huang, L., Feng, Y.-Q. and Geng, Q.-S. (2019) MicroRNA-92b-3p suppresses angiotensin II-induced cardiomyocyte hypertrophy via targeting HAND2. Life Sciences, 232, 116635.

27. Hu, Z.-Q., Luo, J.-F., Yu, X.-J., Zhu, J.-N., Huang, L., Yang, J., Fu, Y.-H., Li, T., Xue, Y.-M., Feng, Y.-Q. et al. (2017) Targeting myocyte-specific enhancer factor 2D contributes to the suppression of cardiac hypertrophic growth by miR-92b-3p in mice. Oncotarget, 8.

28. Chen, Z., Liang, S., Zhao, Y. and Han, Z. (2012) miR-92b regulates Mef2 levels through a negative-feedback circuit during Drosophila muscle development. Development, 139, 3543–3552.

29. Martin, M. (2011) Cutadapt removes adapter sequences from high-throughput sequencing reads. *EMBnet*. journal, 17, 10–12.

30. Chan, P.P. and Lowe, T.M. (2016) GtRNAdb 2.0: an expanded database of transfer RNA genes identified in complete and draft genomes. Nucleic acids research, 44, D184–D189.

31. Ludwig, W., Strunk, O., Westram, R., Richter, L., Meier, H., Yadhukumar, Buchner, A., Lai, T., Steppi, S., Jobb, G., et al. (2004) ARB: a software environment for sequence data. Nucleic Acids Res, 32, 1363–1371.

32. Quast, C., Pruesse, E., Yilmaz, P., Gerken, J., Schweer, T., Yarza, P., Peplies, J. and Glöckner, F.O. (2013) The SILVA ribosomal RNA gene database project: improved data processing and web-based tools. Nucleic acids research, 41, D590–D596.

33. Langmead, B., Trapnell, C., Pop, M. and Salzberg, S.L. (2009) Ultrafast and memory-efficient alignment of short DNA sequences to the human genome. Genome Biology, 10, R25.

34. Friedländer, M.R., Mackowiak, S.D., Li, N., Chen, W. and Rajewsky, N. (2011) miRDeep2 accurately identifies known and hundreds of novel microRNA genes in seven animal clades. Nucleic Acids Research, 40, 37–52.

35. Kozomara, A., Birgaoanu, M. and Griffiths-Jones, S. (2019) miRBase: from microRNA sequences to function. Nucleic Acids Res, 47, D155–d162.

36. Kalvari, I., Nawrocki, E.P., Ontiveros-Palacios, N., Argasinska, J., Lamkiewicz, K., Marz, M., Griffiths-Jones, S., Toffano-Nioche, C., Gautheret, D., Weinberg, Z. et al. (2020) Rfam 14: expanded coverage of metagenomic, viral and microRNA families. Nucleic Acids Research, 49, D192–D200.

37. Liao, Y., Smyth, G.K. and Shi, W. (2014) featureCounts: an efficient general purpose program for assigning sequence reads to genomic features. Bioinformatics, 30, 923–930.

38. Bolger, A.M., Lohse, M. and Usadel, B. (2014) Trimmomatic: a flexible trimmer for Illumina sequence data. Bioinformatics, 30, 2114–2120.

39. Dobin, A., Davis, C.A., Schlesinger, F., Drenkow, J., Zaleski, C., Jha, S., Batut, P., Chaisson, M. and Gingeras, T.R. (2012) STAR: ultrafast universal RNA-seq aligner. Bioinformatics, 29, 15–21.

40. Frankish, A., Diekhans, M., Jungreis, I., Lagarde, J., Loveland, Jane E., Mudge, J.M., Sisu, C., Wright, J.C., Armstrong, J., Barnes, I. et al. (2021) GENCODE 2021. Nucleic Acids Research, 49, D916–D923.

41. Love, M.I., Huber, W. and Anders, S. (2014) Moderated estimation of fold change and dispersion for RNA-seq data with DESeq2. Genome Biology, 15, 550.

42. Floor, S. (2018), GitHub repository.

43. Marco, A. (2018) SeedVicious: Analysis of microRNA target and near-target sites. PLOS ONE, 13, e0195532.

44. Mi, H., Muruganujan, A., Ebert, D., Huang, X. and Thomas, P.D. (2018) PANTHER version 14: more genomes, a new PANTHER GO-slim and improvements in enrichment analysis tools. Nucleic Acids Research, 47, D419–D426.

45. Ru, Y., Kechris, K.J., Tabakoff, B., Hoffman, P., Radcliffe, R.A., Bowler, R., Mahaffey, S., Rossi, S., Calin, G.A., Bemis, L. et al. (2014) The multiMiR R package and database: integration of microRNA-target interactions along with their disease and drug associations. Nucleic acids research, 42, e133–e133.

46. Elliott, D.A., Braam, S.R., Koutsis, K., Ng, E.S., Jenny, R., Lagerqvist, E.L., Biben, C., Hatzistavrou, T., Hirst, C.E., Yu, Q.C. et al. (2011) NKX2-5eGFP/w hESCs for isolation of human cardiac progenitors and cardiomyocytes. Nature Methods, 8, 1037–1040.

47. Giacomelli, E., Meraviglia, V., Campostrini, G., Cochrane, A., Cao, X., van Helden, R.W.J., Krotenberg Garcia, A., Mircea, M., Kostidis, S., Davis, R.P., et al. (2020) Human-iPSC-Derived Cardiac Stromal Cells Enhance Maturation in 3D Cardiac Microtissues and Reveal Non-cardiomyocyte Contributions to Heart Disease. Cell Stem Cell, 26, 862–879.e811.

48. Ng, E.S., Davis, R., Stanley, E.G. and Elefanty, A.G. (2008) A protocol describing the use of a recombinant protein-based, animal product-free medium (APEL) for human embryonic stem cell differentiation as spin embryoid bodies. Nature Protocols, 3, 768–776.

49. Sievers, F. and Higgins, D.G. (2014) Clustal Omega, accurate alignment of very large numbers of sequences. Methods Mol Biol, 1079, 105–116.

50. Ramaiah, M., Tan, K., Plank, T.M., Song, H.W., Dumdie, J.N., Jones, S., Shum, E.Y., Sheridan, S.D., Peterson, K.J., Gromoll, J. et al. (2019) A microRNA cluster in the Fragile-X region expressed during spermatogenesis targets FMR1. EMBO Rep, 20.

51. Wang, Z., Xie, Y., Wang, Y., Morris, D., Wang, S., Oliver, D., Yuan, S., Zayac, K., Bloomquist, S., Zheng, H. et al. (2020) X-linked miR-506 family miRNAs promote FMRP expression in mouse spermatogonia. EMBO reports, 21, e49024.

52. Tsang, J., Zhu, J. and van Oudenaarden, A. (2007) MicroRNA-Mediated Feedback and Feedforward Loops Are Recurrent Network Motifs in Mammals. Molecular Cell, 26, 753–767.

53. Lai, X., Wolkenhauer, O. and Vera, J. (2016) Understanding microRNA-mediated gene regulatory networks through mathematical modelling. Nucleic acids research, 44, 6019–6035.

54. Sharma, A., Wasson, L.K., Willcox, J.A., Morton, S.U., Gorham, J.M., DeLaughter, D.M., Neyazi, M., Schmid, M., Agarwal, R. and Jang, M.Y. (2020) GATA6 mutations in hiPSCs inform mechanisms for maldevelopment of the heart, pancreas, and diaphragm. Elife, 9, e53278.

55. Liufu, Z., Zhao, Y., Guo, L., Miao, G., Xiao, J., Lyu, Y., Chen, Y., Shi, S., Tang, T. and Wu, C.I. (2017) Redundant and incoherent regulations of multiple phenotypes suggest microRNAs’ role in stability control. Genome Res, 27, 1665–1673.

56. Chandradoss, Stanley D., Schirle, Nicole T., Szczepaniak, M., MacRae, Ian J. and Joo, C. (2015) A Dynamic Search Process Underlies MicroRNA Targeting. Cell, 162, 96–107.

57. Hedges, S.B., Dudley, J. and Kumar, S. (2006) TimeTree: a public knowledge-base of divergence times among organisms. Bioinformatics, 22, 2971–2972.

58. Kent, W.J., Sugnet, C.W., Furey, T.S., Roskin, K.M., Pringle, T.H., Zahler, A.M. and Haussler, D. (2002) The human genome browser at UCSC. Genome research, 12, 996–1006.

59. Wilson, K.D., Hu, S., Venkatasubrahmanyam, S., Fu, J.D., Sun, N., Abilez, O.J., Baugh, J.J., Jia, F., Ghosh, Z., Li, R.A. et al. (2010) Dynamic microRNA expression programs during cardiac differentiation of human embryonic stem cells: role for miR-499. Circ Cardiovasc Genet, 3, 426–435.

60. Amin, S., Donaldson, I.J., Zannino, D.A., Hensman, J., Rattray, M., Losa, M., Spitz, F., Ladam, F., Sagerström, C. and Bobola, N. (2015) Hoxa2 selectively enhances Meis binding to change a branchial arch ground state. Developmental cell, 32, 265–277.

61. Donaldson, I.J., Amin, S., Hensman, J.J., Kutejova, E., Rattray, M., Lawrence, N., Hayes, A., Ward, C.M. and Bobola, N. (2012) Genome-wide occupancy links Hoxa2 to Wnt-β-catenin signaling in mouse embryonic development. Nucleic acids research, 40, 3990–4001.

62. Sheehy, N.T., Cordes, K.R., White, M.P., Ivey, K.N. and Srivastava, D. (2010) The neural crest-enriched microRNA miR-452 regulates epithelial-mesenchymal signaling in the first pharyngeal arch. Development, 137, 4307.

63. Takeuchi, J.K., Mileikovskaia, M., Koshiba-Takeuchi, K., Heidt, A.B., Mori, A.D., Arruda, E.P., Gertsenstein, M., Georges, R., Davidson, L., Mo, R. et al. (2005) Tbx20 dose-dependently regulates transcription factor networks required for mouse heart and motoneuron development. Development, 132, 2463–2474.

64. Lepore, J.J., Mericko, P.A., Cheng, L., Lu, M.M., Morrisey, E.E. and Parmacek, M.S. (2006) GATA-6 regulates semaphorin 3C and is required in cardiac neural crest for cardiovascular morphogenesis. The Journal of Clinical Investigation, 116, 929–939.

65. Hoffman, J.I., Kaplan, S. and Liberthson, R.R. (2004) Prevalence of congenital heart disease. American heart journal, 147, 425–439.

66. Ventura, A., Young, A.G., Winslow, M.M., Lintault, L., Meissner, A., Erkeland, S.J., Newman, J., Bronson, R.T., Crowley, D. and Stone, J.R. (2008) Targeted deletion reveals essential and overlapping functions of the miR-17∼ 92 family of miRNA clusters. Cell, 132, 875–886.

67. Zhao, Y., Ransom, J.F., Li, A., Vedantham, V., von Drehle, M., Muth, A.N., Tsuchihashi, T., McManus, M.T., Schwartz, R.J. and Srivastava, D. (2007) Dysregulation of cardiogenesis, cardiac conduction, and cell cycle in mice lacking miRNA-1-2. Cell, 129, 303–317.

68. Goren, Y., Kushnir, M., Zafrir, B., Tabak, S., Lewis, B.S. and Amir, O. (2012) Serum levels of microRNAs in patients with heart failure. European journal of heart failure, 14, 147–154.

69. Martinez, N.J. and Walhout, A.J. (2009) The interplay between transcription factors and microRNAs in genome−scale regulatory networks. Bioessays, 31, 435–445.

70. Cora’, D., Re, A., Caselle, M. and Bussolino, F. (2017) MicroRNA-mediated regulatory circuits: outlook and perspectives. Physical Biology, 14, 045001.

71. Tsuchihashi, T., Maeda, J., Shin, C.H., Ivey, K.N., Black, B.L., Olson, E.N., Yamagishi, H. and Srivastava, D. (2011) Hand2 function in second heart field progenitors is essential for cardiogenesis. Developmental biology, 351, 62–69.

72. Vincentz, J.W., Clouthier, D.E. and Firulli, A.B. (2020) Mis-expression of a cranial neural crest cell-specific gene program in cardiac neural crest cells modulates HAND factor expression, causing cardiac outflow tract phenotypes. Journal of Cardiovascular Development and Disease, 7, 13.

73. Cai, X., Nomura-Kitabayashi, A., Cai, W., Yan, J., Christoffels, V.M. and Cai, C.-L. (2011) Myocardial Tbx20 regulates early atrioventricular canal formation and endocardial epithelial– mesenchymal transition via Bmp2. Developmental biology, 360, 381–390.

74. Laurent, F., Girdziusaite, A., Gamart, J., Barozzi, I., Osterwalder, M., Akiyama, J.A., Lincoln, J., Lopez-Rios, J., Visel, A., Zuniga, A. et al. (2017) HAND2 Target Gene Regulatory Networks Control Atrioventricular Canal and Cardiac Valve Development. Cell Rep, 19, 1602–1613.

75. Shalgi, R., Brosh, R., Oren, M., Pilpel, Y. and Rotter, V. (2009) Coupling transcriptional and post-transcriptional miRNA regulation in the control of cell fate. Aging (Albany NY), 1, 762.

76. Riba, A., Bosia, C., El Baroudi, M., Ollino, L. and Caselle, M. (2014) A combination of transcriptional and microRNA regulation improves the stability of the relative concentrations of target genes. PLoS Comput Biol, 10, e1003490.

77. Alberti, C. and Cochella, L. (2017) A framework for understanding the roles of miRNAs in animal development. Development, 144, 2548–2559.

78. Subasic, D., Brümmer, A., Wu, Y., Pinto, S.M., Imig, J., Keller, M., Jovanovic, M., Lightfoot, H.L., Nasso, S. and Goetze, S. (2015) Cooperative target mRNA destabilization and translation inhibition by miR-58 microRNA family in C. elegans. Genome research, 25, 1680–1691.

79. Marco, A., Hooks, K. and Griffiths-Jones, S. (2012) Evolution and function of the extended miR-2 microRNA family. RNA Biology, 9, 242–248.

80. Chen, J., Huang, Z.-P., Seok, H.Y., Ding, J., Kataoka, M., Zhang, Z., Hu, X., Wang, G., Lin, Z. and Wang, S. (2013) mir-17–92 cluster is required for and sufficient to induce cardiomyocyte proliferation in postnatal and adult hearts. Circulation research, 112, 1557–1566.

81. Danielson, L.S., Park, D.S., Rotllan, N., Chamorro-Jorganes, A., Guijarro, M.V., Fernandez-Hernando, C., Fishman, G.I., Phoon, C.K. and Hernando, E. (2013) Cardiovascular dysregulation of miR−17−92 causes a lethal hypertrophic cardiomyopathy and arrhythmogenesis. The FASEB Journal, 27, 1460–1467.

82. Zhang, X., Azhar, G. and Wei, J.Y. (2012) The expression of microRNA and microRNA clusters in the aging heart. PloS one, 7, e34688.

