## Supplementary Data for "MicroRNA miR-92b-3p regulation of a cardiovascular gene regulatory network in the developing branchial arches"

### Supplementary figure 1

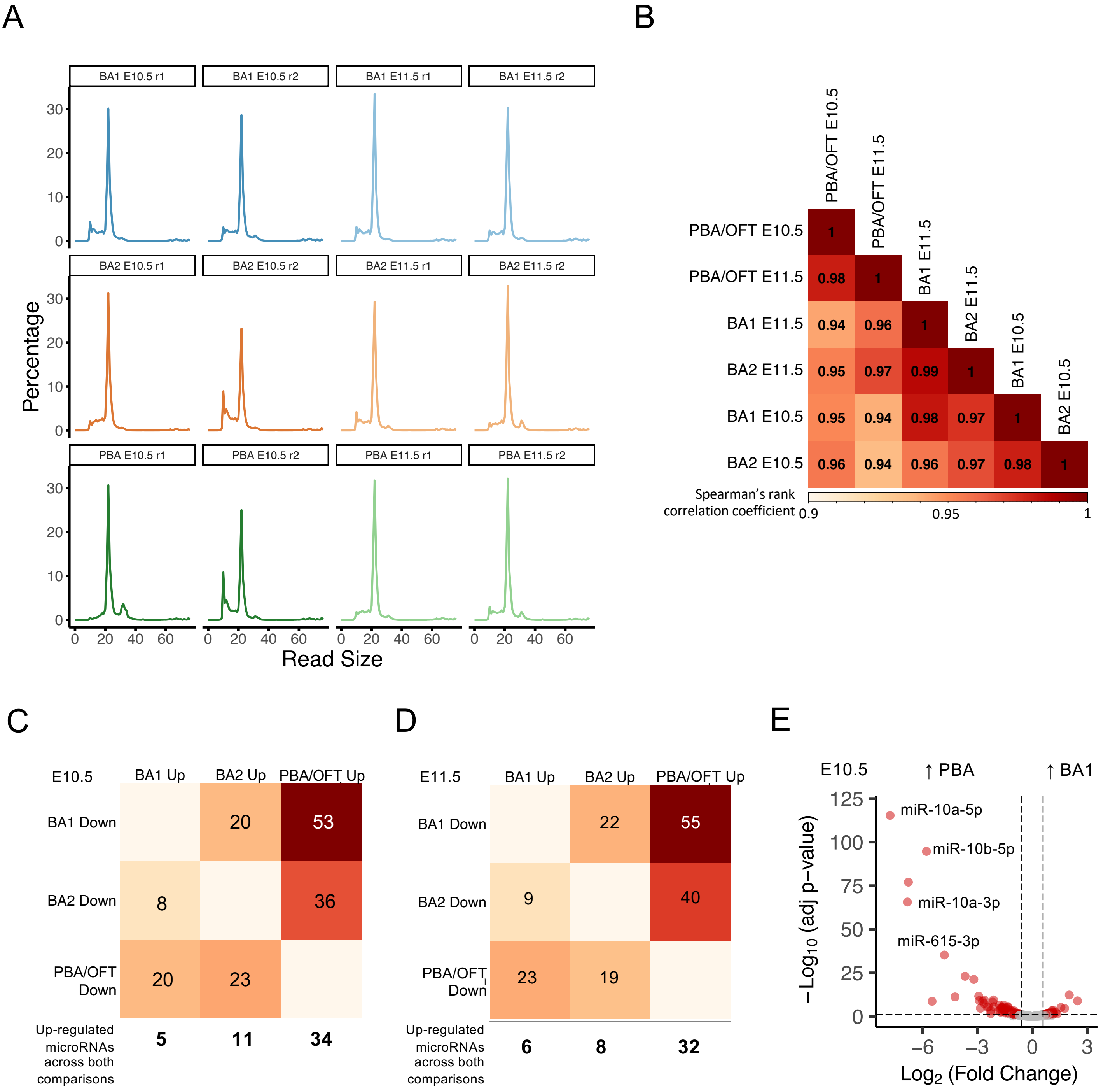


**Supplementary Figure 1. BA small-RNA-seq library features.**

A. Read size across BA small-RNA seq libraries. B. Spearman's rank correlation coefficient calculated using microRNA expression across BA samples. C-D. Number of differentially expressed microRNAs across BA pairwise comparisons at E10.5 and E11.5. E. Differentially expressed microRNAs at E10.5 in BA1 v PBA, with those located nearby PBA/OFT upregulated Hox genes labelled.

### Supplementary figure 2

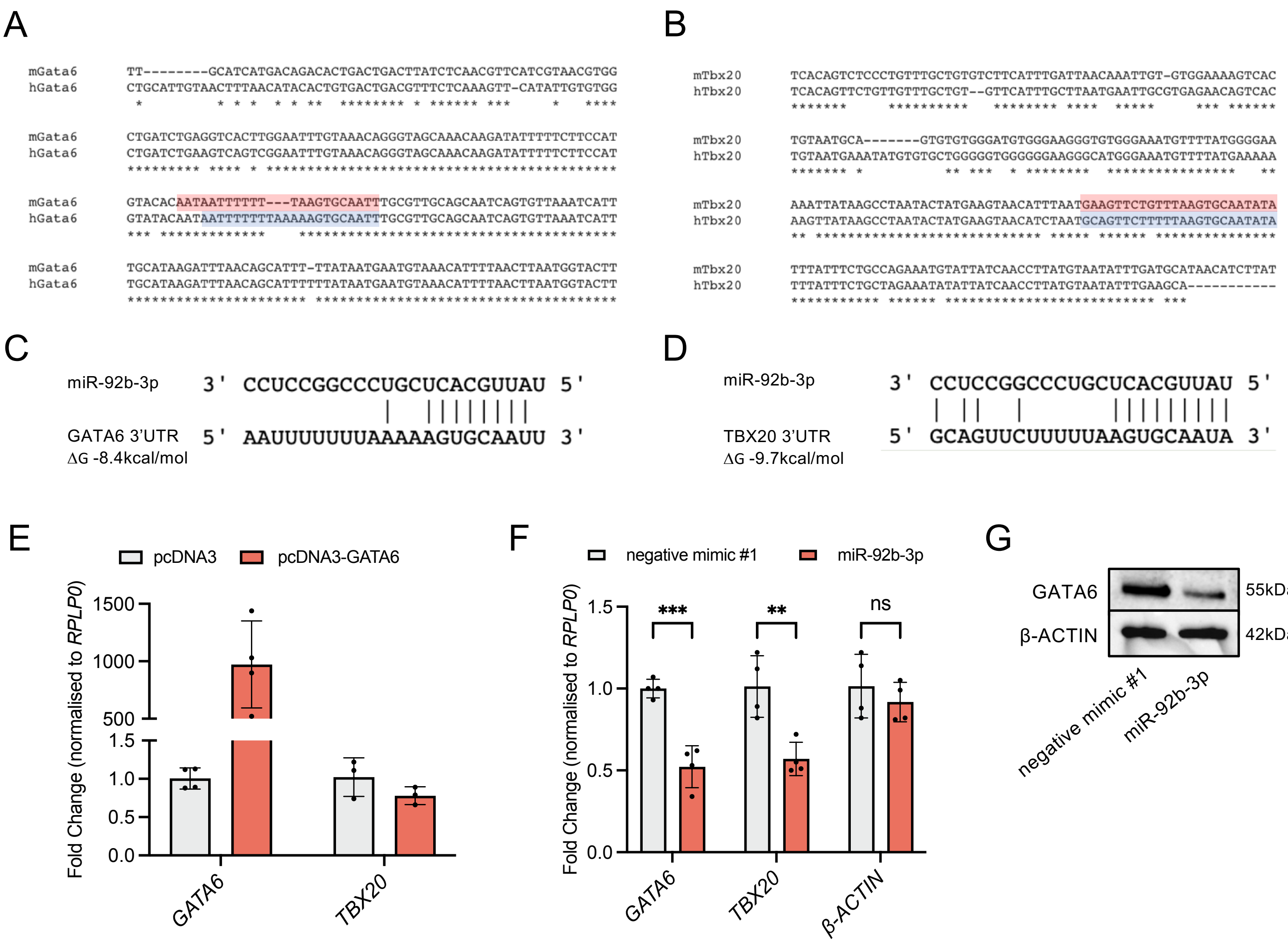


**Supplementary Figure 2. Conservation of miR-92b-3p interaction with *GATA6* and *TBX20*.**

A-B. Alignment between mouse and human *Gata6/GATA6* and *Tbx20/TBX20* 3'UTRs surrounding the region with homologous miR-92b-3p binding sites. C-D. Predicted interactions between miR-92b-3p and *GATA6* and *TBX20* 3'UTRs. E. *GATA6* and *TBX20* in HEK293 cells, 48h after transfection of pcDNA3-GATA6. Samples were first normalised to *RPLP0* and then used to calculate fold change. Values are presented as the mean±s.d., n≥3. F. *GATA6*, *TBX20,* and *ACTB* (negative control not predicted to be targeted by miR-92b-3p), in HEK293 cells 48h after transfection with 30nM microRNA mimics. Samples were normalised to *RPLP0* and then used to calculate fold change. Values are presented as the mean±s.d., n=4. Statistical significance was calculated performing multiple unpaired t-tests, *GATA6* p-value = $4.8 \times 10^{-5}$, *TBX20* p-value = 0.006. G. Western blot of GATA6 and β-ACTIN, following 48h transfection with 30nM microRNA mimics. Replicate western blots showed similar levels of knockdown.

### Supplementary File 1. List of primers

| Primer | Sequence |
|---|---|
| NEBNext Index 1- BA1 E10.5 rep 1 | CAAGCAGAAGACGGCATACGAGATCGTGATGTGACTGGAGTTCAGACGTGTGCTCTTCCGATCT |
| NEBNext Index 2- BA2 E10.5 rep 1 | CAAGCAGAAGACGGCATACGAGATACATCGGTGACTGGAGTTCAGACGTGTGCTCTTCCGATCT |
| NEBNext Index 3- PBA E10.5 rep 1 | CAAGCAGAAGACGGCATACGAGATGCCTAAGTGACTGGAGTTCAGACGTGTGCTCTTCCGATCT |
| NEBNext Index 4- BA1 E10.5 rep 2 | CAAGCAGAAGACGGCATACGAGATTGGTCAGTGACTGGAGTTCAGACGTGTGCTCTTCCGATCT |
| NEBNext Index 5- BA2 E10.5 rep 2 | CAAGCAGAAGACGGCATACGAGATCACTGTGTGACTGGAGTTCAGACGTGTGCTCTTCCGATCT |
| NEBNext Index 6- PBA E10.5 rep 2 | CAAGCAGAAGACGGCATACGAGATATTGGCGTGACTGGAGTTCAGACGTGTGCTCTTCCGATCT |
| NEBNext Index 7- BA1 E11.5 rep 1 | CAAGCAGAAGACGGCATACGAGATGATCTGGTGACTGGAGTTCAGACGTGTGCTCTTCCGATCT |
| NEBNext Index 8- BA2 E11.5 rep 1 | CAAGCAGAAGACGGCATACGAGATTCAAGTGTGACTGGAGTTCAGACGTGTGCTCTTCCGATCT |
| NEBNext Index 9- PBA E11.5 rep 1 | CAAGCAGAAGACGGCATACGAGATCTGATCGTGACTGGAGTTCAGACGTGTGCTCTTCCGATCT |
| NEBNext Index 10- BA1 E11.5 rep 2 | CAAGCAGAAGACGGCATACGAGATAAGCTAGTGACTGGAGTTCAGACGTGTGCTCTTCCGATCT |
| NEBNext Index 11- BA2 E11.5 rep 2 | CAAGCAGAAGACGGCATACGAGATGTAGCCGTGACTGGAGTTCAGACGTGTGCTCTTCCGATCT |
| NEBNext Index 12- PBA E11.5 rep 2 | CAAGCAGAAGACGGCATACGAGATTACAAGGTGACTGGAGTTCAGACGTGTGCTCTTCCGATCT |
| Gata6-3'UTR forward | TAAGCAGAGCTCGAGCTGGTGCTACCAAGAGG |
| Gata6-3'UTR reverse | TAAGCAGTCGACGGTACAGGCGTCAAGAGTGT |
| Gata6-3'UTR mutagenesis forward | GTACACAATAATTTTTTTAAGCGCCCCTTGCGTTGCAGCAATCAGTG |
| Gata6-3'UTR mutagenesis reverse | CACTGATTGCTGCAACGCAAGGGGCGCTTAAAAAAATTATTGTGTAC |
| Tbx20 3'UTR forward | TAAGCAGAGCTCTCACAGTCTCCCTGTTTGCTG |
| Tbx20 3'UTR reverse | TAAGCAGTCGACCAGGTGAGAATATACAGACACCAGA |
| Tbx20 3'UTR mutagenesis forward | GTTTAAGTGCCCCATATTTATTTCTGCCAGAAATGTATTATCAAC |
| Tbx20 3'UTR mutagenesis reverse | GTTGATAATACATTTCTGGCAGAAATAAATATGGGGCACTTAAAC |
| Hand1-3'UTR forward | TAAGCAGAGCTCGCGGTTGCAACTACCCACTA |
| Hand1-3'UTR reverse | TAAGCAGTCGACCAGCAACGAATGGGAACGC |
| GATA6 RT-qPCR forward | CCACAACACAACCTACAGCC |
| GATA6 RT-qPCR reverse | ACGCCTATGTAGAGCCCATC |
| TBX20 RT-qPCR forward | GAGGGAAAGTGTGGAGAGCC |
| TBX20 RT-qPCR reverse | AAGGCTGACCCTCGATTTGG |
| HAND1 RT-qPCR forward | TCTTCCACCCTTTTGGAGCG |
| HAND1 RT-qPCR reverse | GCCTTTCATCTTCCTGCGTC |
| RPLP0 RT-qPCR forward | CACCATTGAAATCCTGAGTGATGT |
| RPLP0 RT-qPCR reverse | TGACCAGCCCAAAGGAGAAG |
| ZNF503 RT-qPCR forward | GCACGAACTGGCCACATTTT |
| ZNF503 RT-qPCR reverse | GGGCTCTTCTTGGCATCGAG |
| ACTB RT-qPCR forward | GGCTGTATTCCCCTCCATCG |
| ACTB RT-qPCR reverse | CCAGTTGGTAACAATGCCATG |
| pmirGLO sequencing primer forward | GGCAAGATCGCCGTGTAATTC |
| pmirGLO sequencing primer reverse | TATCATGTCTGCTCGAAGCG |

#### Supplementary File 2. High-confidence novel mouse microRNAs predicted using miRDeep2 (Friedländer et al., 2011)

| miRDeep2 microRNA ID | Shared seed with mouse | Mature sequence [1] | Star sequence [2] | Genomic location | Relative location |
|---|---|---|---|---|---|
| chr1_2334 | - | uguguguaaggguccugucaau | ugugcagguccuguacauucagug | chr1:106669442..106669504:- | *Bcl20*, intron |
| chr2_6424 | - | agcagcagcuggagcagugggggc | ccucuccuccggcggccgcggc | chr2:153444424..153444481:- | *Nol4l*, intron |
| chr2_5073 | - | ucaaggucuagcagguagcacagc | gugagugccugcugcccugaguu | chr2:181379689..181379758:+ | *Zgpat*, exon |
| chr2_3301 | - | uauucugacucuaccuaacugga | ucugcuagguaguggugugaacugc | chr2:18177241..18177305:+ | *Mllt10*, intron |
| chr3_9116 | - | cugagccccgagacuguauuac | uuacaguccgagcccugagacu | chr3:121770700..121770760:- | *Abcd3*, intron |
| chr5_14235 | - | uaugaguucuaggcauugaaucu | gucaaugcucugaacuccuaag | chr5:33618090..33618153:- | *Fam53a*, intron |
| chr7_20796 | - | ugacucucugcuuccccccagu | uuaggggguugcugagggaccu | chr7:126375202..126375257:- | *Spns1*, intron, <= 10kb miR-7058 |
| chr8_23232 | - | uguccgggggaccgacuuugcc | auggagccgcgcuccgggacgcgu | chr8:105496918..105496972:- | intergenic |
| chr8_22037 | - | ucagccguucagcaccguccagaca | acugggcaaggaacaggcucggg | chr8:105660297..105660358:+ | *Ctc*f, intron |
| chr8_22488 | - | aaacugucugucuguucauagc | cauaugagcagaccuccaguuu | chr8:13885880..13885935:- | *Coprs,* intron |
| chr9_24954 | *mmu-miR-1839-5p* | uagguagaccaggcugaucu | auccuccugucucugccuucu | chr9:122981876..122981935:+ | *Kif15*, intron |
| chr9_23920 | - | ccaggcugcuggagucugggu | ccagucuccaucugccuuccu | chr9:40696225..40696280:+ | *Clmp*, intron |
| chr9_25372 | - | ucagacugcugagucacauugc | aaugugacucagcuaccugaac | chr9:52103707..52103764:- | *Gm27686*, exon |
| chr9_24142 | *mmu-miR-702-5p* | gugaguggagacucgguagaggu | ccucuccuguucuccaucuagu | chr9:57141915..57141995:+ | *Man2c1*, intron |
| chr9_25805 | - | aaaagaacugcguggcuaucuggc | ugagguagucagcagugaucuuu | chr9:94520299..94520363:- | *Dip2ka*, 3'UTR |
| chr10_26210 | - | gagggacauacucaaugagaac | ucuuauuguccaugucccugcc | chr10:4092813..4092871:+ | *Mthfd1l*, intron |
| chr11_30225 | - | uccugcccccuuucccuguaga | uaaggguggauggggcagagcu | chr11:105347182..105347235:+ | *Mrc2*, intron |
| chr11_28554 | - | uggcugccagcagagccuggau | ccacucugcugggcagcccag | chr11:4675148..4675209:+ | *Ascc2*, intron |
| chr11_30626 | - | ucuguuggauccugugaggaca | cccauaggauuuuaacagaacu | chr11:5080868..5080921:- | *Ewsr1*, intron |
| chr11_31180 | - | uagcguucucggagaucauga | ucugaugacugagauugcugacc | chr11:61546572..61546631:- | *Epn2*, exon |
| chr11_30658 | - | cuugaucuucucccucugcagg | ugcggaaggacagauccugugg | chr11:6264032..6264090:- | *Ddx56*, intron |
| chr12_33597 | - | ucaagugugacaagaucucuac | ugaguuaucuugaggcacuugac | chr12:13245067..13245126:- | *Ddx1*, intron |
| chr12_32939 | - | ugcugaauccagagguuacaucu | uggugaucucggagauucagg | chr12:73284657..73284720:+ | *Trmt5*, antisense exon |
| chr15_39753 | - | aauagcuugggacaucagcucugu | agaacugaugacugagcaagg | chr15:36595713..36595768:- | *Pabpc1*, 3'UTR |
| chr16_41858 | - | ucaugugucucuuguguugauc | ccggcacaacagaaacaugagu | chr16:44490355..44490412:- | *Boc*, intron |
| chr16_41185 | - | aacauguuggacggugcacu | augcgcugucagucaacauggc | chr16:64858038..64858103:+ | *Cggbp1*, 3'UTR |
| chr19_47164 | - | ugucuugggcucuggaguugaguu | aucgcugcugagccuagacugg | chr19:45773947..45774009:- | *Oga*, intron |
| chr19_46558 | - | ugggcuccgccuguggucgc | cugacuuccaggcccagcccugca | chr19:57467284..57467344:+ | *Trub1*, intron |

**Supplementary File 3. miR-92b-3p PBA/OFT subset target predictions.**

| miRNA | gene_name | position | type | value | 3’UTRlength | seed |
|---|---|---|---|---|---|---|
| MIMAT0004899 | ENSMUSG00000000631 | 711 | 7_m8 | -7.4 | 1148 | AUUGCA |
| MIMAT0004899 | ENSMUSG00000004872 | 1365 | 8mer | -11.1 | 2041 | AUUGCA |
| MIMAT0004899 | ENSMUSG00000005836 | 665 | 7_m8 | -9.5 | 1156 | AUUGCA |
| MIMAT0004899 | ENSMUSG00000008136 | 304 | 7_m8 | -9.5 | 443 | AUUGCA |
| MIMAT0004899 | ENSMUSG00000021109 | 546 | 7_A1 | -9.2 | 1830 | AUUGCA |
| MIMAT0004899 | ENSMUSG00000022443 | 579 | 7_m8 | -10.7 | 1315 | AUUGCA |
| MIMAT0004899 | ENSMUSG00000022803 | 483 | 7_A1 | -8.2 | 608 | AUUGCA |
| MIMAT0004899 | ENSMUSG00000024529 | 1074 | 7_m8 | -7.8 | 3163 | AUUGCA |
| MIMAT0004899 | ENSMUSG00000024593 | 3397 | 7_A1 | -9.3 | 3415 | AUUGCA |
| MIMAT0004899 | ENSMUSG00000025809 | 999 | 7_m8 | -7.64 | 2529 | AUUGCA |
| MIMAT0004899 | ENSMUSG00000025809 | 1080 | 7_m8 | -12 | 2529 | AUUGCA |
| MIMAT0004899 | ENSMUSG00000025880 | 1387 | 7_m8 | -8 | 1573 | AUUGCA |
| MIMAT0004899 | ENSMUSG00000026185 | 3674 | 7_m8 | -8.6 | 4448 | AUUGCA |
| MIMAT0004899 | ENSMUSG00000027474 | 1267 | 7_A1 | -7.2 | 1643 | AUUGCA |
| MIMAT0004899 | ENSMUSG00000027887 | 2905 | 7_A1 | -8.1 | 3067 | AUUGCA |
| MIMAT0004899 | ENSMUSG00000030790 | 299 | 7_m8 | -9.4 | 655 | AUUGCA |
| MIMAT0004899 | ENSMUSG00000031965 | 4538 | 8mer | -9.7 | 7317 | AUUGCA |
| MIMAT0004899 | ENSMUSG00000034460 | 784 | 7_A1 | -8.1 | 5173 | AUUGCA |
| MIMAT0004899 | ENSMUSG00000036867 | 209 | 7_m8 | -9.3 | 451 | AUUGCA |
| MIMAT0004899 | ENSMUSG00000037335 | 184 | 7_m8 | -7.2 | 897 | AUUGCA |
| MIMAT0004899 | ENSMUSG00000038193 | 128 | 7_m8 | -9.87 | 776 | AUUGCA |
| MIMAT0004899 | ENSMUSG00000040118 | 1078 | 7_m8 | -9.3 | 3840 | AUUGCA |
| MIMAT0004899 | ENSMUSG00000041842 | 899 | 7_m8 | -7.59 | 2272 | AUUGCA |
| MIMAT0004899 | ENSMUSG00000042942 | 1650 | 8mer | -10.84 | 2460 | AUUGCA |
| MIMAT0004899 | ENSMUSG00000044447 | 782 | 8mer | -9.1 | 4286 | AUUGCA |
| MIMAT0004899 | ENSMUSG00000045092 | 1248 | 7_m8 | -10.19 | 1365 | AUUGCA |
| MIMAT0004899 | ENSMUSG00000049281 | 1555 | 8mer | -7.8 | 3184 | AUUGCA |
| MIMAT0004899 | ENSMUSG00000052374 | 41 | 7_A1 | -9.6 | 71 | AUUGCA |
| MIMAT0004899 | ENSMUSG00000055022 | 1375 | 7_m8 | -14.9 | 2381 | AUUGCA |
| MIMAT0004899 | ENSMUSG00000062991 | 901 | 7_m8 | -8.3 | 3480 | AUUGCA |
| MIMAT0004899 | ENSMUSG00000063632 | 1386 | 8mer | -7.74 | 6960 | AUUGCA |
